# *Vangl* facilitates mesenchymal thinning during lung sacculation independently of *Celsr*

**DOI:** 10.1101/2022.12.28.522148

**Authors:** Sarah V. Paramore, Carolina Trenado-Yuste, Rishabh Sharan, Danelle Devenport, Celeste M. Nelson

**Affiliations:** Departments of Molecular Biology, Princeton University, Princeton, NJ 08544; Departments of Chemical & Biological Engineering, Princeton University, Princeton, NJ 08544; Departments of Lewis-Sigler Institute for Integrative Genomics, Princeton University, Princeton, NJ 08544

**Keywords:** tissue morphodynamics, mechanical force, motility

## Abstract

The planar cell polarity (PCP) complex orients cytoskeletal and multicellular organization throughout vertebrate development. PCP is speculated to function in formation of the murine lung, where branching morphogenesis generates a complex tree of tubular epithelia whose distal tips expand dramatically during sacculation in preparation for gas exchange after birth. Here, using tissue-specific knockouts, we show that the PCP complex is dispensable in the airway epithelium for sacculation. Rather, we find a novel, *Celsr1*-independent role for the PCP component *Vangl* in the pulmonary mesenchyme: loss of *Vangl1/2* inhibits mesenchymal thinning and expansion of the saccular epithelium. Further, loss of mesenchymal *Wnt5a* mimics the sacculation defects observed in *Vangl2*-mutant lungs, implicating mesenchymal Wnt5a/Vangl signaling as a key regulator of late lung morphogenesis. By mathematically modeling sacculation, we predict that the process of sacculation requires a fluid mesenchymal compartment. Finally, lineage-tracing and cell-shape analyses are consistent with the pulmonary mesenchyme acting as a fluid tissue, and suggest that loss of *Vangl1/2* likely impacts the ability of mesenchymal cells to exchange neighbors. Our data thus uncover an explicit function for *Vangl* and the pulmonary mesenchyme during late lung morphogenesis to actively shape the saccular epithelium.

## Introduction

During development of the mammalian lung, the pulmonary epithelium undergoes dozens of rounds of stereotyped branching to form a complex airway tree (Metzger et al., 2008). After branching is completed, the lungs undergo the canalicular and saccular stages of development. The pulmonary mesenchyme thins while distal airways widen dramatically to increase epithelial surface area, forming saccules. Concurrently, the airway epithelium differentiates into specialized cell types including type I and type II alveolar cells, secretory cells, and ciliated cells, preparing the lung to function after birth (Desai et al., 2014; Morrisey and Hogan, 2010). Understanding the processes that drive sacculation is essential for motivating new strategies to treat defects associated with prematurity and congenital abnormalities.

During sacculation, the mesenchyme between distal airways thins to a roughly single-cell layer as the epithelium expands. This rapid change in tissue shape results in an increase in epithelial surface area that supports gas exchange postnatally. While the specific cellular mechanisms that underlie epithelial expansion and mesenchymal thinning are unknown, decades of studies have made clear that mechanical forces generated by lumenal fluid pressure are an important driver of sacculation. For example, fetal breathing movements, which begin at approximately *E*16 in mice, lead to cyclic increases in fluid pressure and promote sacculation (Liggins et al., 1981; Moessinger et al., 1990; Wigglesworth and Desai, 1982). Conversely, sacculation is halted when the pressure in the lung is reduced by leakage of amniotic fluid (Perlman et al., 1976) or by ablation of fetal breathing movements via loss of skeletal muscle (Tseng et al., 2000), leading to lung hypoplasia. Pressure of the fluid within the lung has a direct effect on the fate of alveolar epithelial cells: those exposed to high pressure differentiate into type I alveolar epithelial cells (AEC1s) whereas those exposed to low pressure differentiate into type II alveolar epithelial cells (AEC2s) (Li et al., 2018). However, it is unknown if pressure also regulates mesenchymal thinning. One could posit that a high lumenal fluid pressure may push mesenchymal cells into a thin layer, as increasing the pressure of this fluid increases the overall rate of lung development (Nelson et al., 2017; Unbekandt et al., 2008). Mesenchymal cells might also play an active role in rearranging both themselves and the surrounding extracellular matrix (ECM) to facilitate saccular expansion.

The core planar cell polarity (PCP) complex was recently implicated in the process of sacculation (Poobalasingam et al., 2017; Yates et al., 2010). The core PCP complex consists of three transmembrane proteins [in mice, Celsr1-3, Vangl1/2, and Frizzled3/6 (Wang et al., 2006)] that localize asymmetrically along the plane of a tissue and regulate oriented cell behaviors (Devenport, 2014). To generate asymmetry and relay polarity information, Vangl/Celsr localize to the opposite side of the cell as Frizzled/Celsr. The Celsr cadherin repeats mediate the formation of PCP junctions between neighboring cells (**Fig. 1A**). This asymmetric localization is indicative of and required for PCP function. In vertebrates, PCP regulates cytoskeletal organization, and is required for convergent extension movements during gastrulation and neural tube closure (Curtin et al., 2003; Heisenberg et al., 2000; Tada and Smith, 2000; Torban et al., 2004; Wallingford et al., 2000; Wang et al., 2006). Similarly, PCP regulates the shape of epithelial tubules in the developing kidney via convergent extension (Kunimoto et al., 2017; Lienkamp et al., 2012). In the lung, the role of PCP is beginning to be appreciated: at the cellular level, this complex regulates the orientation of airway cilia (Vladar et al., 2012), and sacculation and alveologenesis are perturbed in PCP-mutant lungs (Poobalasingam et al., 2017; Yates et al., 2010). However, it is unclear why loss of PCP causes defects in late-stage lung development. Thus, the idea that the PCP complex regulates epithelial widening during sacculation remains an attractive hypothesis.

**Figure 1.**
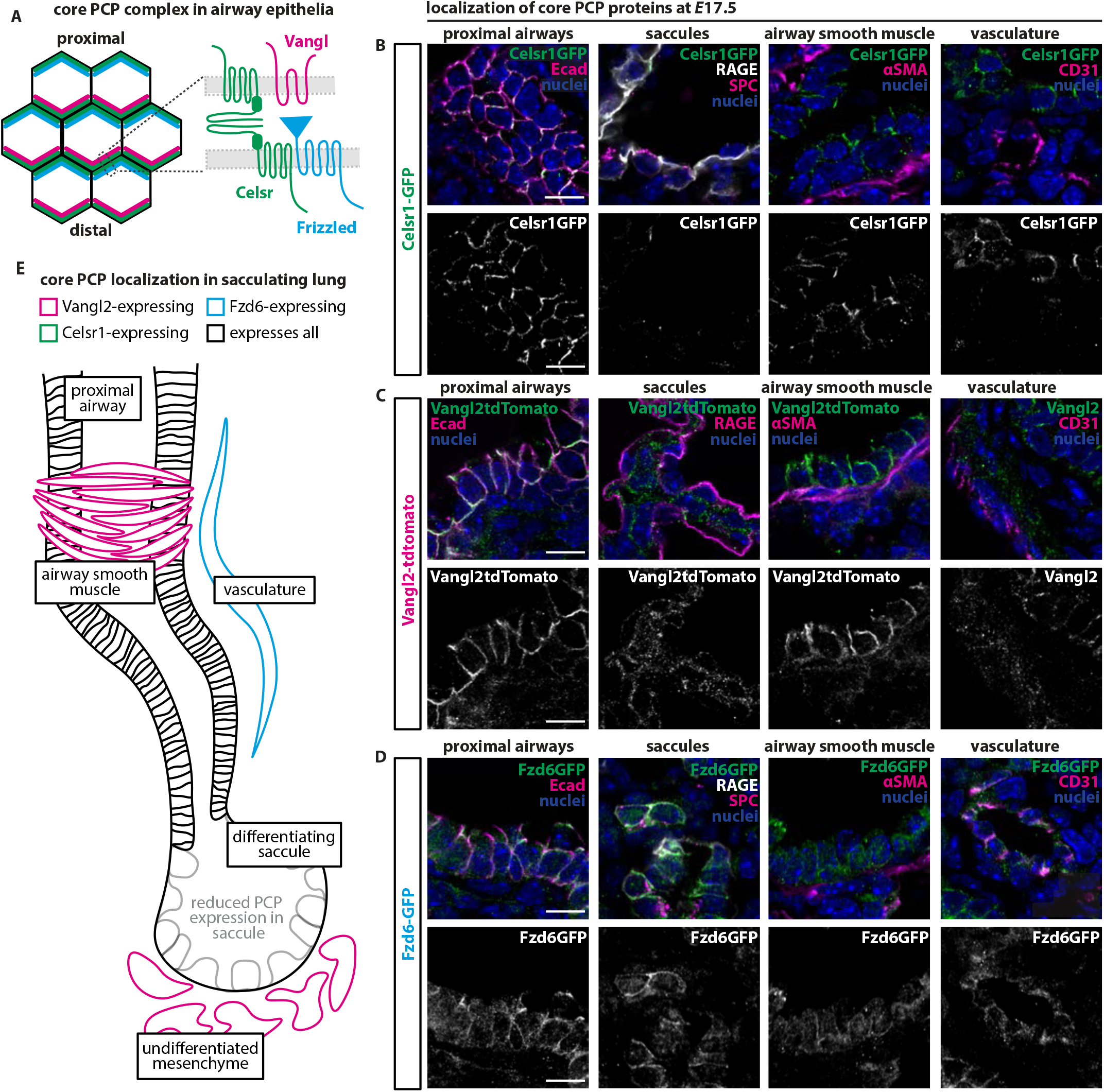
Core PCP proteins are differentially expressed in sacculation-stage lungs. **A**, Schematic illustrating the PCP complex in a planar epithelial tissue. **B**, Representative images of 10-μm-thick sections of *E*17.5 lungs showing localization of Celsr1-GFP in proximal airways (Ecad), saccules (RAGE and SPC), airway smooth muscle (αSMA), and vasculature (CD31). **C**, Representative images of 10-μm-thick sections of *E*17.5 lungs showing localization of Vangl2-tdTomato or Vangl2 in proximal airways (Ecad), saccules (RAGE and SPC), airway smooth muscle (αSMA), and vasculature (CD31). **D**, Representative images of 10-μm-thick sections of *E*17.5 lungs showing localization of Fzd6-GFP in proximal airways (Ecad), saccules (RAGE and SPC), airway smooth muscle (αSMA), and vasculature (CD31). All scale bars, 10 μm. **E**, Schematic summarizing localization of core PCP proteins in the sacculation-stage lung.

Here, we investigated the cellular mechanisms through which the PCP complex influences sacculation. We reveal that *Celsr1* is expressed only in the lung epithelium and mesothelium, while *Vangl2* is expressed ubiquitously throughout the lung, including in the pulmonary mesenchyme. We find severe sacculation defects in the lungs of *Vangl2-*mutants, but not *Celsr1-*mutants. Surprisingly, this phenotype is not reproduced when *Vangl1* and *Vangl2* are depleted from the lung epithelium; instead, sacculation defects arise from loss of *Vangl1/2* in the lung mesenchyme. We also show that loss of mesenchymal *Wnt5a* mimics the sacculation defects observed in *Vangl2*-mutant lungs, implicating *Wnt5a/Vangl* signaling in late-stage lung development. Using a mathematical model of sacculation, we predict that increasing the fluidity of the mesenchyme increases the extent of sacculation. Lineage-labeling experiments and cell-shape analyses suggest that mesenchymal cells actively exchange neighbors during sacculation, and these cellular behaviors are likely perturbed in *Vangl*-mutants. Our data thus reveal that the pulmonary mesenchyme plays a key, active role in shaping the architecture of the distal lung in a *Vangl*-dependent, *Celsr*-independent manner. These findings add to a growing body of evidence suggesting that PCP genes, whose functions are best understood in planar epithelia, also play important roles mesenchymal tissues to facilitate morphogenesis (Kunimoto et al., 2017; Rao-Bhatia et al., 2020; Zhang et al., 2020).

## Materials and Methods

### Mouse lines and breeding

All procedures involving animals were approved by Princeton University’s Institutional Animal Care and Use Committee (IACUC). Mice were housed in an AAALAC-accredited facility in accordance with the NIH Guide for the Care and Use of Laboratory Animals. This study was compliant with all relevant ethical regulations regarding animal research. *Celsr1GFP, Fzd6GFP* and *Vangl2tdTomato* mouse lines were used to assess the localization of core PCP proteins during lung development (Basta et al., 2021). *Vangl2*^*Lp/Lp*^ embryos (Kibar et al., 2001) and *Celsr1*^*Crsh/Crsh*^ embryos (Curtin et al., 2003) were used to determine how loss of PCP function affects sacculation. *ShhCreGFP; Vangl1*^*fl/fl*^; *Vangl2* ^*fl/fl*^; *Rosa26mTmG/Rosa26mTmG* embryos were used to conditionally delete *Vangl1/2* from the lung epithelium (Copley et al., 2013; Harfe et al., 2004; Harris et al., 2006). *Dermo1Cre; Vangl1* ^*fl/fl*^; *Vangl2* ^*fl/fl*^; *Rosa26mTmG/Rosa26mTmG* embryos were used to conditionally delete *Vangl1/2* from the pulmonary mesenchyme (Yin et al., 2008; Yu et al., 2003). *Tbx4-rtTA; Tet-O-Cre; Vangl1* ^*fl/fl*^; *Vangl2* ^*fl/fl*^; *Rosa26mTmG/Rosa26mTmG* embryos were used to conditionally delete *Vangl1/2* from the pulmonary mesenchyme in an inducible manner (Zhang et al., 2013). To induce full *Vangl1/2* deletion, both doxycycline-medicated water (0.5 mg/mL) and an intraperitoneal injection (0.1 mg doxycycline/1 g weight) were administered to pregnant dams at *E*14. *Tbx4-rtTA; Tet-O-Cre; R26R-Confetti* (Livet et al., 2007) embryos were used for lineage-labeling and cell-shape analysis. *Tbx4-rtTA; Tet-O-Cre; Vangl1* ^*fl/fl*^; *Vangl2* ^*fl/fl*^; *R26R-Confetti* embryos were used for cell-shape analysis. *ShhCreGFP; Wnt5a* ^*fl/fl*^ embryos were used to conditionally delete *Wnt5a* from the lung epithelium (Ryu et al., 2013). *Tbx4-rtTA; TetO-Cre; Wnt5a* ^*fl/fl*^ embryos were used to conditionally delete *Wnt5a* from the pulmonary mesenchyme in an inducible manner; doxycycline was administered in water at *E*15 as described above. Genotyping primers are detailed in **Supplementary Methods**.

### Immunofluorescence analysis

Lungs and tracheas were dissected from embryos in PBS and fixed in 4% paraformaldehyde (PFA). All tracheas were fixed for 1 h at 4°C, washed with PBS, incubated overnight in blocking buffer, washed with PBS, incubated with primary antibody overnight, washed with PBS, and incubated with secondary antibody overnight. Tracheas were then washed and mounted on a slide for imaging in Prolong Gold. *E*13.5 lungs were fixed for 30 min at 4°C. *E*16-18.5 lungs were fixed for 1 h at 4°C. Lungs were washed in PBS and taken through a sucrose gradient before embedding in OCT. 10-μm-thick and 200-μm-thick frozen sections were obtained from samples using a Leica CM3050S cryostat. 10-μm-thick sections were washed in PBT3 (PBS with 0.3% Triton X-100), washed in PBS, incubated in blocking buffer for 1 h at room temperature, and then incubated in blocking buffer with primary antibody overnight. Slides were then washed in PBS and incubated with secondary antibody for 3 h and mounted in Prolong Gold. 200-μm-thick floating sections were washed in PBT3, washed in PBS, incubated in blocking buffer overnight at 4°C, and then incubated in blocking buffer with primary antibody overnight. Sections were then washed in PBS and incubated with secondary antibody overnight at 4°C. Sections were then washed in PBS and mounted in Prolong Gold. Antibodies used for staining are detailed in **Supplementary Methods**.

### Quantification of Celsr1 polarity

Celsr1 polarity was calculated using Packing Analyzer V2 software (Aigouy et al., 2010) as previously described (Aw et al., 2016). The software measures the axis and magnitude (nematic order) of junctional polarity. Cells were segmented using the E-cadherin signal. The angles were plotted in a circular histogram using the polar plot function in MATLAB. The magnitude of polarity and orientation of average Celsr1 polarity were overlaid as a line on top of the histogram, where the length of the line and its orientation reflect the magnitude and direction of Celsr1 polarity.

### Single-cell RNA-sequencing (scRNA-seq) analysis

We analyzed a published scRNA-seq dataset (GSE149563) (Zepp et al., 2021) generated from *E*17.5 mouse lungs. This analysis was carried out using the Seurat package (Butler et al., 2018). The data were first filtered to exclude cells with fewer than 500 genes, more than 30,000 unique molecular identifiers (possible multiplets), and greater than 10% mitochondrial DNA (dying cells). Following the Seurat pipeline, we then normalized the data, identified variable features, scaled gene expression for each cell, ran principal components analysis, and identified neighbors and clusters. We then generated uniform manifold approximation and projection (UMAP) plots and extracted cluster markers to identify cell types. Clusters from the *E*17.5 lung dataset representing distinct cell types were annotated based on the most highly expressed genes in each cluster. We then examined the expression patterns of genes of interest by color-coding the UMAP and comparing cell-level expression of these genes in different cell clusters.

### Sacculation analysis

Tiled 20× images of entire lung sections were acquired on a Nikon A1RSi confocal microscope. Sections were imaged for RAGE, SPC, and Hoechst, or for mTmG, RAGE, and Hoechst when applicable. To calculate the area of tissue, the image was first binarized such that non-tissue regions were labeled in white. After binarization, this image was negated and the sum of the white (tissue) pixels was calculated, giving the value of the area of the tissue.

To calculate the area and perimeter of each saccule from a given image, the image was first binarized where the saccules were labeled in white. The area and perimeter were then calculated for each saccule using the ‘regionprops’ function in MATLAB. This value was then converted to units of μm^2^. Any saccule smaller than 50 μm^2^ (for images with no transgenic mTmG) or 25 μm^2^ (for images with transgenic mTmG) was filtered out and discarded from further analysis.

### Computational model of sacculation

To test the role of mesenchymal fluidity during sacculation, we created a 2D agent-based model of particles (cells) undergoing Brownian motion, as described by their center of mass. The model combines epithelial and mesenchymal compartments in which cells are reduced to their center points and packed such that they do not overlap. The motion of a cell in both compartments is affected by its interactions with neighboring cells and fluctuations in the system. The equations of motion for the positions of cells in both compartments are considered in the overdamped limit, which is commonly used to describe biological systems where the inertial effects are much smaller than the effects from intracellular interactions or the fluctuations in the system.

We modeled the mesenchyme as a collection of cells, where each cell is reduced to its center and is driven by a Langevin-like equation in continuous-time. The equation of motion for a mesenchymal cell *i* is described as follows:

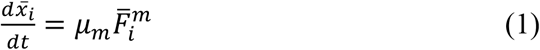

where 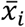 is the position of the cell *i, μ*_*m*_ is the mobility coefficient for the mesenchymal population, and 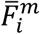 is the sum of all forces acting upon the *i-th* mesenchymal cell:

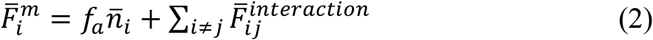

The term 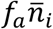 models the internal cellular processes that drive a mesenchymal cell to move in the direction of its polarity, and the interaction between the *i-th* cell and its nearest neighbors (Fumoto et al., 2017; Fumoto et al., 2019; Higaki et al., 2017; Li et al., 2018; Takigawa-Imamura et al., 2015) is defined as:

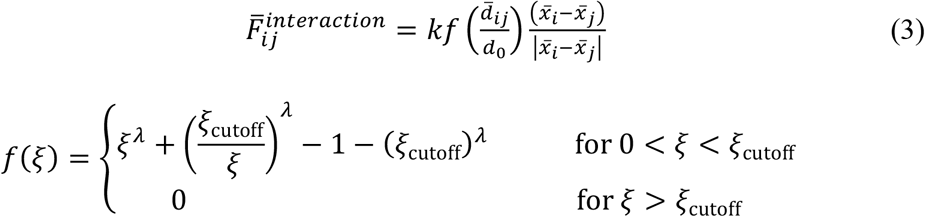

where 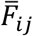 is the force between *i*-*th* and *j*-*th* mesenchymal cells (repulsive at short distances and attractive at longer distances), and *k* describes the strength of the interaction force, and 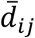 is the distance between the cells calculated as 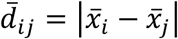. The optimal value *d*_0_ = *r*_*i*_ + *r*_*j*_ is the optimal distance between two cells. To simplify the model, we fixed the optimal distance to *d*_0_ = 1 (See **Figure 5A**). Cell-cell interactions are also assumed to occur indirectly between the mesenchyme and the epithelium using the same force defined in **Equation 3**, so that the deformation of the epithelial tissue is coupled to that of the surrounding mesenchyme. The parameters are *ξ*_cutoff_ =1.63, *λ* = 2, and *k* =1.

The polarity vector 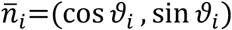 is modeled as a Gaussian white rotational noise,

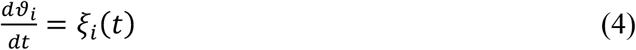

where *ξ*_*i*_(*t*) is a white noise with zero mean ⟨*ξ*_*i*_(*t*)⟩ = *ξ* and correlations ⟨*ξ*_*i*_(*t*)*ξ*_*j*_(*t*′)⟩ = 2*D*_*r*_*δ*_*i,j*_*δ*(*t* − *t*′). *D*_*r*_ measures the average magnitude of the stochastic force and is a rotational diffusion coefficient.

We modeled the structure of the epithelium as a 2D chain of cells distributed in a single layer that initially conformed to the circumference of a saccule. The equation of motion for an epithelial cell *i* is described as:

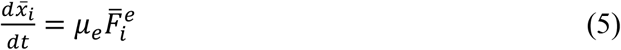

where *μ*_*e*_ is the mobility coefficient for the epithelium, and 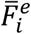 denote the summations of the forces acting upon the *i-th* epithelial cell

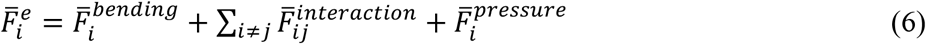

The forces are considered to mimic the deformation of the epithelium observed in the experiments. The effect of bending is incorporated to align the cells and obtain a smooth outline of the epithelium (Fumoto et al., 2017) as follows

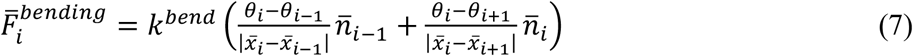

where *θ*_*i*_ represents the angle between the vectors 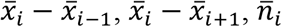 is a unit vector normal to 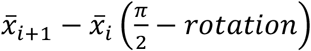, and *k*^*bend*^ is the bending elasticity coefficient. Epithelial cell interactions were modeled using **Equation 3**.

Expansion of the epithelium in response to lumenal fluid pressure (Fumoto et al., 2017; Fumoto et al., 2019; Higaki et al., 2017) was modeled by incorporating an outward force on the epithelial cells as

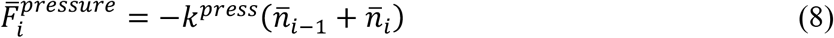

where *k*^*press*^ represents its magnitude.

Although the output of the simulations represents the epithelium as a continuous line, the simulated tissue is comprised of discrete cells whose interactions are within a distance *ξ*_cutoff_ =1.63 (See pink curve **Figure 5A**). *In vivo*, epithelial surface area increases because of cell-shape changes; to simplify the model, the increase in epithelial surface area *in silico* is accounted for by increased epithelial cell division instead. Cell proliferation was implemented computationally by measuring the distance between two adjacent cells. If the distance is larger than d = 0.5, one cell is added in between two pre-existing epithelial cells. This approach allowed us to model the growth and deformation of the epithelium.

### Lineage-tracing analysis

For each cell of each clone (simulated and *in vivo)*, the center point was determined by identifying the xy coordinates using FIJI. This was then graphed using a custom MATLAB script where, for each clone, the clone centroid was calculated by the mean of each point. The distance from the center of each cell to its clone centroid was calculated as the Euclidean distance. Plots were then generated where each center point of each cell of a clone was subtracted by the centroid of that clone. These clone-centroid subtracted points were then overlayed on top of each other and colored by the embryonic stage.

### Mesenchymal cell-shape analysis

Mesenchymal cells were segmented and binarized using the ilastik 2-stage autocontext workflow (Berg et al., 2019), using the signal from cytoplasmic RFP^+^ mesenchymal cells. Segmented images were then analyzed in MATLAB by labeling each connected component using the bwlabel function. The per segmented cell metrics of perimeter, area, convex hull area, circularity, and major and minor axis lengths were then calculated using the regionprops function. Area/convex hull area was calculated as the ratio between area and convex hull area. Aspect ratio was calculated as the ratio of the major and minor axis lengths. Shape factor was calculated as the ratio of the perimeter and the square root of the area. Histograms were then generated using R and the ggplot2 package (Team, 2021; Villanueva and Chen, 2019).

## Results

### Core PCP components have different expression patterns during sacculation

To investigate the expression and possible role for PCP components during sacculation, we harvested *E*17.5 lungs from transgenic mice in which endogenous core PCP components are fluorescently tagged, allowing us to assess the localization of Celsr1, Vangl2, and Fzd6 (**Fig. 1A**) (Basta et al., 2021). We investigated PCP expression across multiple tissues within the lung, including the proximal airways, developing saccules, airway smooth muscle, and vasculature. As expected, Celsr1-GFP localizes predominantly to the airway epithelium and is absent from mesenchymal tissues and endothelium (**Fig. 1B**). However, the fluorescence intensity of Celsr1-GFP is reduced in the saccular epithelium, suggesting that the expression of Celsr1 might be downregulated in these cells at this stage of development (**Fig. 1B**). Similarly, Vangl2 is highly expressed in the proximal epithelium, with reduced expression in saccules (**Fig. 1C**). Surprisingly, we observed essentially ubiquitous expression of Vangl2 in the pulmonary mesenchyme, where it is membrane-localized in both airway smooth muscle and undifferentiated mesenchyme adjacent to saccules (**Fig. 1C**). Fzd6 is expressed at high levels in proximal epithelium and endothelial cells, but appears to be excluded from airway smooth muscle and saccule-adjacent mesenchyme (**Fig. 1D**).

To assess the expression of the other components of the core PCP complex, we analyzed existing scRNA-seq data from *E*17.5 lungs (Zepp et al., 2021) (**Supp. Fig. 1A**). Like *Celsr1, Celsr2* is only expressed in the epithelium, whereas *Celsr3* is completely absent from the lung at this stage of development (**Supp. Fig. 1B**). *Vangl1* is expressed in the same cell populations as *Vangl2*, albeit to a lesser extent (**Supp. Fig. 1C**). Finally, *Fzd3* and *Fzd6* are expressed in a small fraction of mesenchymal cells, with higher expression in endothelial, hematopoetic, and epithelial cells (**Supp. Fig. 1D**). These data reveal that while all core transmembrane PCP components are expressed in the airway epithelium, *Vangl1/2* are enriched in mesenchymal populations in which *Celsr1-3* and *Fzd3/6* are largely absent. Further, *Fzd3/6* are expressed in the pulmonary endothelium independent of *Vangl1/2* and *Celsr1-3*. This compartment-specific expression of different core PCP genes suggests that PCP components may have roles outside of the core PCP complex.

### Vangl2^Lp/Lp^ but not Celsr1^Crsh/Crsh^ lungs fail to undergo sacculation

To begin to characterize the role of PCP in the formation of saccules, we harvested lungs at *E*18.5 when sacculation is mostly complete. At this stage of development, PCP-mutants have complete craniorachischisis and severe axis-elongation defects (Curtin et al., 2003; Torban et al., 2004). To distinguish between direct effects from loss of PCP and indirect effects from the open neural tube (ONT) and axis-elongation defects, we took advantage of the incomplete penetrance of the ONT phenotype in our *Celsr1*^*Crsh/Crsh*^ line. Specifically, we compared lungs from ONT *Celsr1*^*Crsh/Crsh*^ embryos to those that presented with a closed neural tube (CNT), which display no defects in axis elongation (unpublished observations). In wild-type lungs, PCP proteins including Celsr1 become polarized in the tracheal epithelium beginning at *E*14.5 (Vladar et al., 2012).

Despite the CNT phenotype of the *Celsr1*^*Crsh/Crsh*^ embryos, there is a clear lack of Celsr1 polarization in the tracheas of these animals, indicating loss of PCP function (**Supp. Fig. 2A-B**). Analysis of *E*18.5 ONT *Celsr1*^*Crsh/Crsh*^ embryos revealed two phenotypically distinct populations: ∼40% of the embryos were pink and had blood flow despite the ONT, while the other ∼60% were white and appeared to be either dead or dying (**Supp. Fig. 2C**). Lung tissue from white *E*18.5 ONT embryos appeared highly abnormal by immunofluorescence (**Supp. Fig. 2D**), and so we excluded these samples from our subsequent analysis of ONT *Celsr1*^*Crsh/Crsh*^ embryos (**Supp. Fig. 2E-F**).

**Figure 2.**
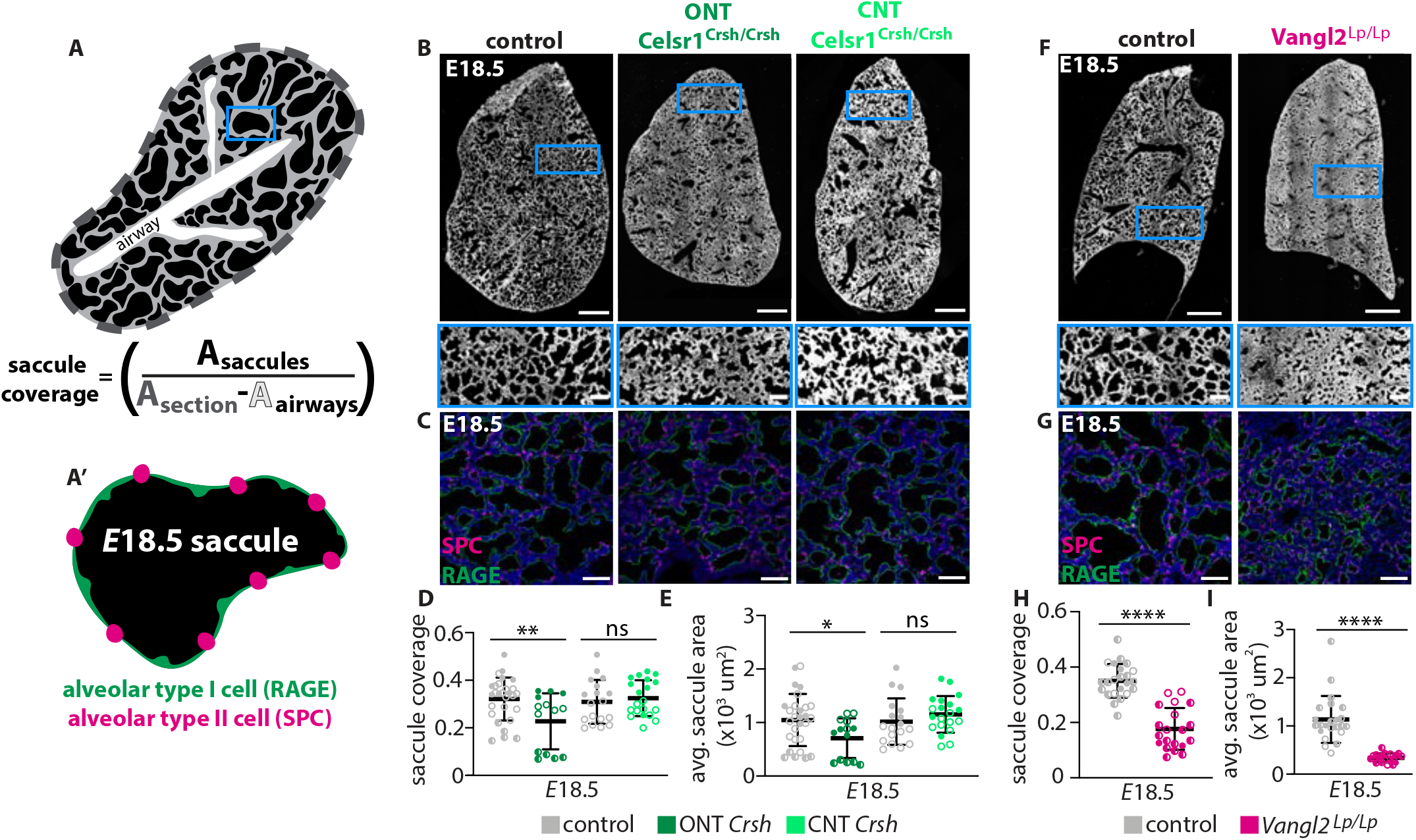
Sacculation fails in *Vangl2*^*Lp/Lp*^ but not *Celsr1*^*Crsh/Crsh*^ embryos. **A**, Schematic illustrating metric used to quantify saccule coverage, where A denotes area; **A’**, Illustration of an *E*18.5 saccule and the marker proteins used to identify AEC1s and AEC2s. **B**, Representative tiled images (scale bars, 500 μm) and insets (scale bars, 100 μm) of *E*18.5 lung sections from control, ONT, and CNT *Celsr1*^*Crsh/Crsh*^ lungs; fluorescence intensity includes signal from staining for SPC, RAGE, and Hoechst. **C**, Representative images of sections of distal lung tissue at *E*18.5 in control, ONT *Celsr1*^*Crsh/Crsh*^, and CNT *Celsr1*^*Crsh/Crsh*^ lungs; scale bars, 50 μm. **D**, Quantification of the percentage of lung area accounted for by saccules at *E*18.5 in control, ONT *Celsr1*^*Crsh/Crsh*^ (*n*=6 control and *n*=3 mutant lungs, *p*=0.0066 via unpaired Student’s t-test), and CNT *Celsr1*^*Crsh/Crsh*^ lungs (*n*=4 control and *n*=4 mutant lungs, *p*=0.5795 via unpaired Student’s t-test); each data point is one lung lobe. **E**, Quantification of average saccular lumenal area at *E*18.5 in control, ONT *Celsr1*^*Crsh/Crsh*^ (*n*=6 control and *n*=3 mutant lungs, *p*=0.0271 via unpaired Student’s t-test), and CNT *Celsr1*^*Crsh/Crsh*^ lungs (*n*=4 control and *n*=4 mutant lungs, *p*=0.2918 via unpaired Student’s t-test); each data point is one lung lobe. **F**, Representative tiled images (scale bars, 500 μm) and insets (scale bars, 100 μm) of *E*18.5 lung sections from control and *Vangl2*^*Lp/Lp*^ lungs; fluorescence intensity includes signal from staining for SPC, RAGE, and Hoechst. **G**, Representative images of sections of distal lung tissue at *E*18.5 in control and *Vangl2*^*Lp/Lp*^ lungs; scale bars, 50 μm. **H**, Quantification of the percentage of lung area accounted for by saccules at *E*18.5 in control and *Vangl2*^*Lp/Lp*^ lungs (*n*=6 control and *n*=5 mutant lungs, *p*<0.0001 via unpaired Student’s t-test); each data point is one lung lobe. **I**, Quantification of average saccular lumenal area at *E*18.5 in control and *Vangl2*^*Lp/Lp*^ lungs (*n*=6 control and *n*=5 mutant lungs, *p*<0.0001 via unpaired Student’s t-test); each data point is one lung lobe. Shown are mean ± s.d; * p < 0.05; ** p < 0.01; **** p < 0.0001. In all graphs, different shapes represent distinct experimental replicates.

To quantify the extent of sacculation, we acquired tiled, confocal images of whole lung sections, which enabled us to assess an entire lobe rather than a small sampling of images. We generated a custom sacculation-analysis pipeline to measure both the area of individual saccules and the fractional area of the lung section accounted for by saccules, which we termed “saccule coverage” (**Fig. 2A**). We found that both the saccule coverage and average saccule area are slightly reduced in lungs from ONT *Celsr1*^*Crsh/Crsh*^ embryos compared to controls at *E*18.5 (**Fig. 2B-E**). However, we observed no differences between lungs from CNT *Celsr1*^*Crsh/Crsh*^ embryos and controls (**Fig. 2B-E**). Furthermore, CNT *Celsr1*^*Crsh/Crsh*^ pups are viable and can breathe after birth, consistent with these animals achieving normal sacculation. These data suggest that the slight reduction in sacculation observed in the ONT *Celsr1*^*Crsh/Crsh*^ embryos is a consequence of gross abnormalities of the embryo, rather than due to loss of *Celsr1* in the lung per se. In contrast, *Vangl2*^*Lp/Lp*^ lungs exhibit severe sacculation defects (**Fig. 2F-G**). At *E*18.5, *Vangl2*^*Lp/Lp*^ lungs have significantly reduced saccule coverage as compared to controls (**Fig. 2H**) and significantly smaller saccule areas (**Fig. 2I**). Thus, our data reveal a role for *Vangl*, but not *Celsr*, in regulating sacculation of the embryonic mouse lung.

### Vangl1/2 is required in the mesenchyme for sacculation

Because *Vangl1/2* are expressed in both mesenchymal and epithelial compartments during sacculation, we took advantage of tissue-specific knockouts to determine whether sacculation requires *Vangl1/2* in the epithelium, the mesenchyme, or both. We generated *ShhCre*; *Vangl1*^*fl/fl*^; *Vangl2*^*fl/fl*^ (epiCKO) embryos, which have conditional deletion of *Vangl1/2* in the airway epithelium and show loss of PCP asymmetry (**Supp. Fig. 3A-C**). We also generated *Dermo1Cre*; *Vangl1*^*fl/fl*^; *Vangl2*^*fl/fl*^ (mesCKO) embryos, in which broad expression of *Cre* in the splanchnic mesoderm leads to conditional deletion of *Vangl1/2* from the pulmonary mesenchyme but not the epithelium (**Supp. Fig. 3D**) (Yin et al., 2008; Yu et al., 2003). Of note, the *Dermo1Cre* allele and the *Vangl2* gene both reside on Chromosome 1, so generating *Dermo1Cre; Vangl2*^*fl/fl*^ embryos required meiotic recombination between the Vangl2 locus and the *Dermo1Cre* insertion site. As expected, mesenchymal loss of *Vangl1/2* does not affect Celsr1 polarity in the tracheal epithelium (**Supp. Fig. 3E-F**).

**Figure 3.**
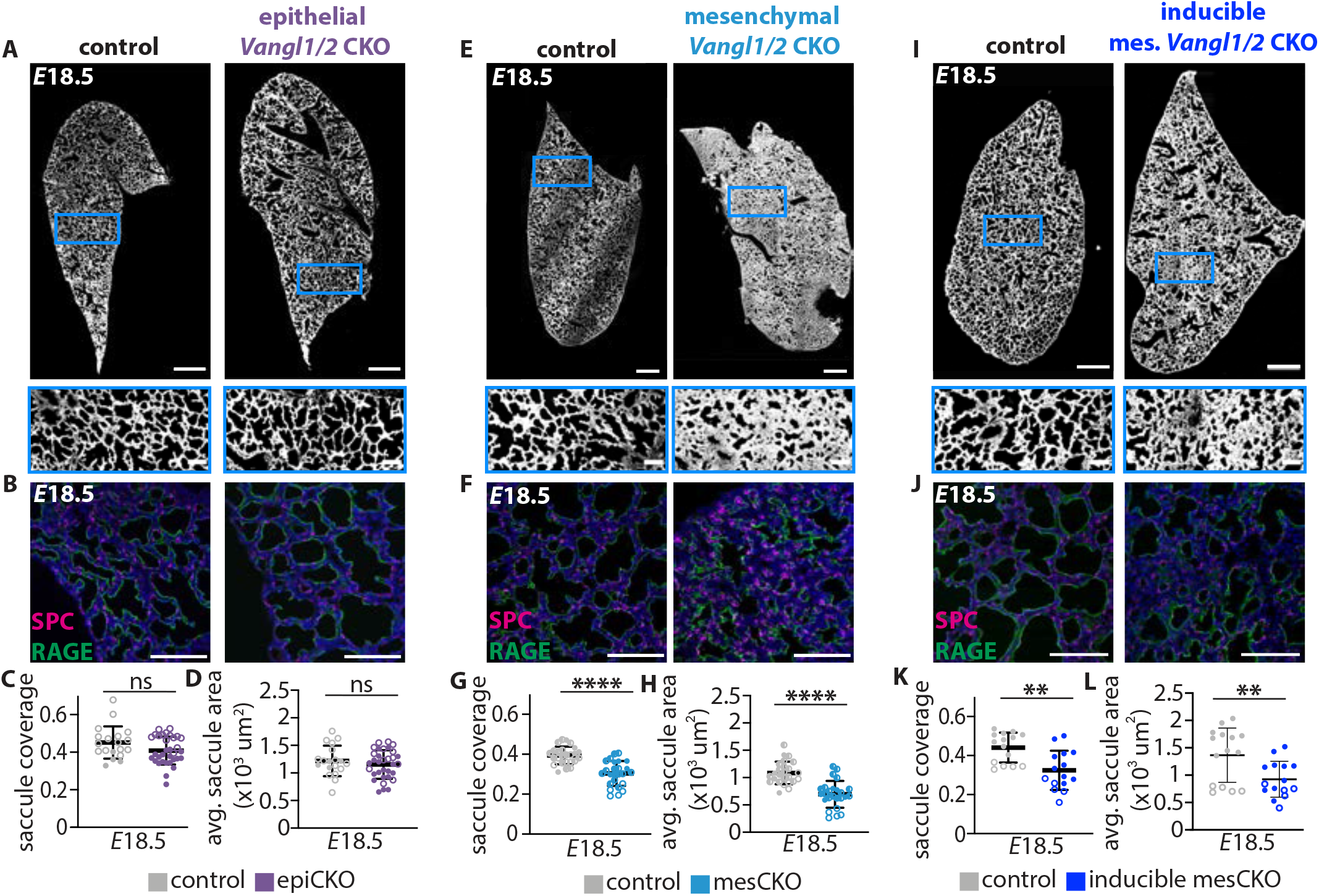
Sacculation requires *Vangl1/2* in the pulmonary mesenchyme but not the airway epithelium. **A**, Representative tiled images (scale bars, 500 μm) and insets (scale bars, 100 μm) of *E*18.5 lung sections from control and epiCKO lungs; fluorescence intensity includes signal from staining for RAGE and Hoechst and transgenic mTmG fluorescence. **B**, Representative images of sections of distal lung tissue at *E*18.5 in control and epiCKO lungs; scale bars, 100 μm. **C**, Quantification of the percentage of lung area accounted for by saccules at *E*18.5 in control and epiCKO lungs (*n*=4 control and *n*=7 mutant lungs, *p*=0.0620 via unpaired Student’s t-test); each data point represents one lung lobe. **D**, Quantification of average saccular lumenal area at *E*18.5 in control and epiCKO lungs (*n*=4 control and *n*=7 mutant lungs, *p*=0.4020 via unpaired Student’s t-test); each data point represents one lung lobe. **E**, Representative tiled images (scale bars, 500 μm) and insets (scale bars, 100 μm) of *E*18.5 lung sections from control and mesCKO lungs; fluorescence intensity includes signal from staining for RAGE and Hoechst and transgenic mTmG fluorescence. **F**, Representative images of sections of distal lung tissue at *E*18.5 in control and mesCKO lungs; scale bars, 100 μm. **G**, Quantification of the percentage of lung area accounted for by saccules at *E*18.5 in control and mesCKO lungs (*n*=5 control and *n*=5 mutant lungs, *p*<0.0001 via unpaired Student’s t-test); each data point represents one lung lobe. **H**, Quantification of average saccular lumenal area at *E*18.5 in control and mesCKO lungs (*n*=5 control and *n*=5 mutant lungs, *p*<0.0001 via unpaired Student’s t-test); each data point represents one lung lobe. **I**, Representative tiled images (scale bars, 500 μm) and insets (scale bars, 100 μm) of *E*18.5 lung sections from control and inducible mesCKO lungs; fluorescence intensity includes staining for RAGE and Hoechst and transgenic mTmG fluorescence. **J**, Representative images of sections of distal lung tissue at *E*18.5 in control and inducible mesCKO lungs; scale bars, 100 μm. **K**, Quantification of the percentage of lung area accounted for by saccules at *E*18.5 in control and inducible mesCKO lungs (*n*=3 control and *n*=3 mutant lungs, *p*=0.0013 via unpaired Student’s t-test); each data point represents one lung lobe. **L**, Quantification of average saccular lumenal area at *E*18.5 in control and inducible mesCKO lungs (*n*=3 control and *n*=3 mutant lungs, *p*=0.0076 via unpaired Student’s t-test); each data point represents one lung lobe. Shown are mean ± s.d; ** p < 0.01; **** p < 0.0001. In all graphs, different shapes represent distinct experimental replicates.

We found that epiCKO lungs undergo sacculation and are indistinguishable from controls at *E*18.5. (**Fig. 3A-D**). In contrast, mesCKO lungs exhibit sacculation defects similar to those observed in *Vangl2*^*Lp/Lp*^ lungs, with significantly reduced saccule coverage, smaller saccular areas, and thicker mesenchyme (**Fig. 3E-H**). To confirm that *Vangl1/2* are specifically required in the pulmonary mesenchyme during sacculation, we generated a second mesenchymal knockout line with an inducible *Cre* (*Tbx4-rtTA; Tet-O-Cre; Vangl1*^*fl/fl*^; *Vangl2*^*fl/fl*^; inducible mesCKO), in which *Cre* expression is restricted to the pulmonary mesenchyme (Zhang et al., 2013). We introduced doxycycline via drinking water and intraperitoneal injection at *E*14.5 to ensure loss of Vangl1/2 protein by *E*16.5, as Vangl2 can remain stable at the membrane for more than a day (unpublished observations). We found that inducible mesCKO lungs have sacculation defects at *E*18.5, phenocopying both the *Vangl2*^*Lp/Lp*^ and mesCKO animals (**Fig. 3I-L**). Notably, if a single functional allele of *Vangl1* or *Vangl2* remained in the embryo (i.e., *Vangl1*^*fl/+*^; *Vangl2*^*fl/fl*^ or *Vangl1*^*fl/fl*^; *Vangl2*^*fl/+*^) no defects were observed and mice were viable postnatally; thus, *Vangl1* and *Vangl2* compensate for each other in this tissue. To determine whether the sacculation defects are caused by changes in cell proliferation or apoptosis, we performed immunofluorescence analysis for phospho-histone-3 and cleaved caspase-3 on control and mesCKO lungs at *E*16.5-18.5. We found no differences in proliferation or apoptosis between controls and mutants (**Supp. Fig. 3G-H**). Altogether, these data suggest that the sacculation defects observed in *Vangl2*^*Lp/Lp*^ embryos are a consequence of a loss of *Vangl* function specifically in the pulmonary mesenchyme during sacculation, rather than due to a requirement for *Vangl* or the PCP complex in the lung epithelium.

### Loss of Wnt5a in the pulmonary mesenchyme mimics loss of mesenchymal Vangl1/2

In epithelia, Vangl2 functions in concert with Celsr1 to affect diverse outcomes ranging from convergent extension to oriented ciliogenesis (Boutin et al., 2014; Cetera et al., 2017; Devenport and Fuchs, 2008; Nishimura et al., 2012; Shi et al., 2014; Stahley et al., 2021; Usami et al., 2021; Vladar et al., 2012). In the pulmonary mesenchyme, however, our data show that Vangl2 acts independently of Celsr1 to promote sacculation. Outside of its core PCP-binding partners, *Vangl2* has been reported to physically or genetically interact with non-canonical Wnt receptor tyrosine kinases, including *Ror2, Ptk7*, and *Ryk* (Andre et al., 2012; Gao et al., 2011; Lu et al., 2004). Analysis of scRNA-seq data from *E*17.5 lungs (Zepp et al., 2021) revealed that *Ror2, Ptk7*, and *Ryk* are all expressed in the mesenchyme (**Fig. 4A, Supp. Fig. 1A**). Of these receptors, *Ryk* shows the strongest expression in the lung during this developmental period, with highest expression in the lung mesenchyme. Each of these receptors has been hypothesized or shown to function downstream of Wnt5a, a non-canonical Wnt ligand (Gao et al., 2011; Martinez et al., 2015; Wang et al., 2020). Consistently, the scRNA-seq analysis also shows that *Wnt5a* is expressed predominantly in the lung mesenchyme at *E*17.5 and at lower levels in the epithelium (**Fig. 4A**). In many developmental contexts, including in tracheal morphogenesis and alveologenesis, a *Wnt5a/Ror2/Vangl* signaling pathway has been implicated in mesenchymal cell polarization and migration (Gao et al., 2011; Kishimoto et al., 2018; Li et al., 2020; Zhang et al., 2020). Given that embryo-wide loss of *Wnt5a* immediately prior to sacculation phenocopies the defects observed in our *Vangl1/2* mesCKO lungs (Li et al., 2020), we hypothesized *Wnt5a* may be upstream of *Vangl* function in the sacculating lung.

**Figure 4.**
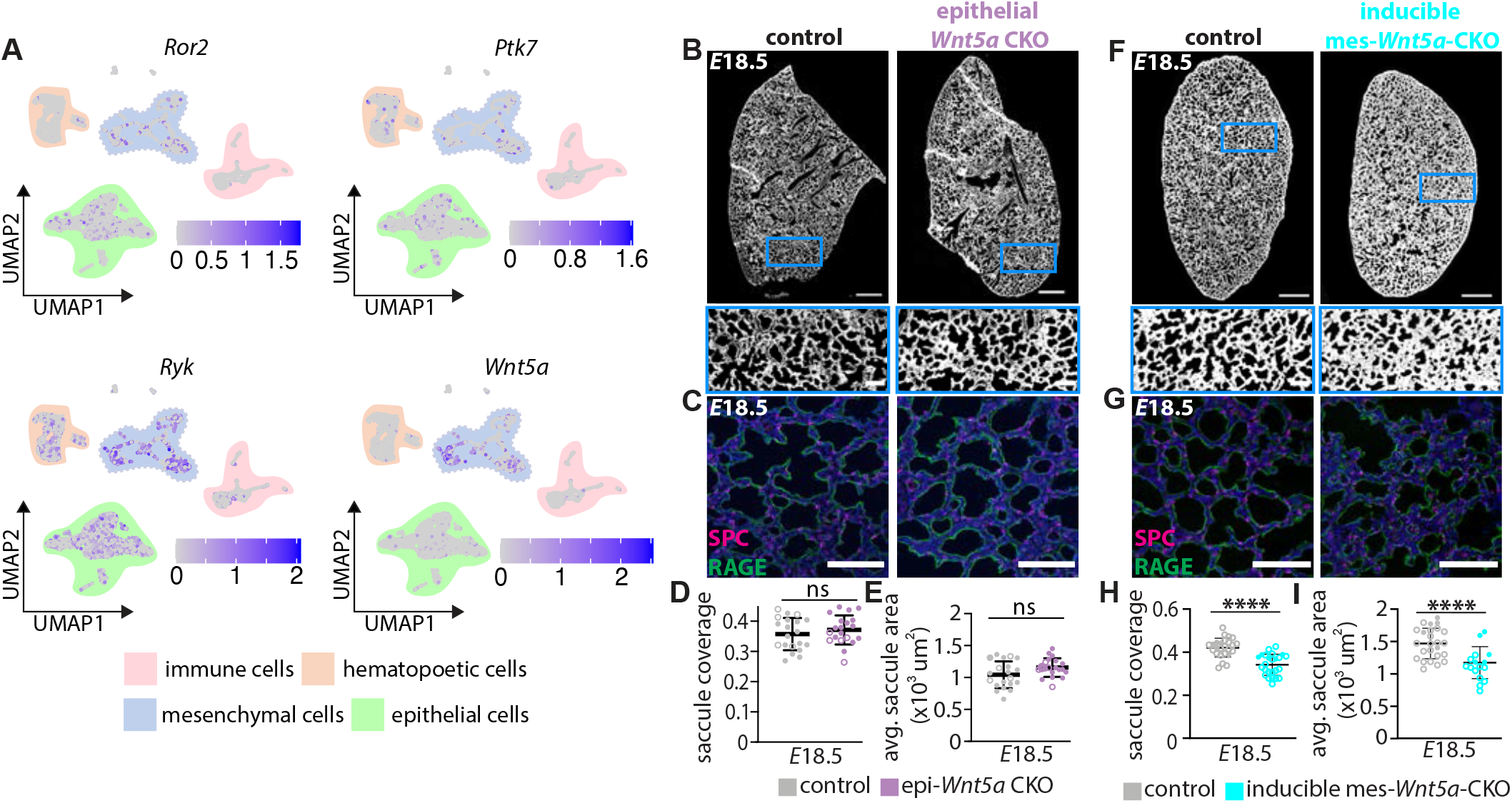
Loss of mesenchymal *Wnt5a* mimics loss of *Vangl1/2* during sacculation. **A**, scRNA-seq analysis of *E*17.5 lungs, data from (Zepp et al., 2021). Cells are clustered into four main groups: immune, epithelial, hematopoetic, and mesenchymal cells. UMAP projections reveal the relative expression of *Ror2, Ptk7, Ryk*, and *Wnt5a*. **B**, Representative tiled images (scale bars, 500 μm) and insets (scale bars, 100 μm) of *E*18.5 lung sections from control and epi-*Wnt5a*-CKO lungs; fluorescence intensity includes signal from staining for SPC, RAGE, and Hoechst. **C**, Representative images of sections of distal lung tissue at *E*18.5 in control and epi-*Wnt5a*-CKO lungs; scale bars, 100 μm. **D**, Quantification of the percentage of lung area accounted for by saccules at *E*18.5 in control and epi-*Wnt5a*-CKO lungs (*n*=4 control and *n*=4 mutant lungs, *p*=0.3994 via unpaired Student’s t-test); each data point represents one lung lobe. **E**, Quantification of average saccular lumenal area at *E*18.5 in control and epi-*Wnt5a*-CKO lungs (*n*=4 control and *n*=4 mutant lungs, *p*=0.0552 via unpaired Student’s t-test); each data point represents one lung lobe. **F**, Representative tiled images (scale bars, 500 μm) and insets (scale bars, 100 μm) of *E*18.5 lung sections from control and inducible mes-*Wnt5a*-CKO lungs; fluorescence intensity includes signal from staining for SPC, RAGE and Hoechst. **G**, Representative images of sections of distal lung tissue at *E*18.5 in control and inducible mes-*Wnt5a*-CKO lungs; scale bars, 100 μm. **H**, Quantification of the percentage of lung area accounted for by saccules at *E*18.5 in inducible mes-*Wnt5a*-CKO lungs (*n*=5 control and *n*=5 mutant lungs, *p* < 0.0001 via unpaired Student’s t-test); each data point represents one lung lobe. **I**, Quantification of average saccular lumenal area at *E*18.5 in control and inducible mes-*Wnt5a*-CKO lungs (*n*=5 control and *n*=5 mutant lungs, *p* < 0.0001 via unpaired Student’s t-test); each data point represents one lung lobe. Shown are mean ± s.d; **** p < 0.0001. In all graphs, different shapes represent distinct experimental replicates.

To determine whether Wnt5a could play a role in Vangl1/2-mediated epithelial expansion and mesenchymal thinning, we generated epithelial (*ShhCre; Wnt5a*^*fl/fl*^) and inducible-mesenchymal (*Tbx4-rtTA; TetOCre; Wnt5a*^*fl/fl*^) *Wnt5a* knockout embryos. Surprisingly, we found that loss of *Wnt5a* in the lung epithelium has no effect on sacculation (**Fig. 4B-E**). In contrast, inducible deletion of *Wnt5a* from the pulmonary mesenchyme at *E*15.5 results in sacculation defects that phenocopy loss of mesenchymal *Vangl1/2* (**Fig. 4F-I**). Mesenchymal expression of *Wnt5a* and *Vangl1/2* are thus required for sacculation of the lung.

### Mathematical modeling predicts a role for mesenchymal cell rearrangements during sacculation

Our observation that Vangl is required in the mesenchyme for sacculation was surprising, as the nonplanar organization of mesenchymal cells is inconsistent with any obvious axis of planar polarization. As a part of its role in the PCP complex, Vangl promotes the asymmetric localization of cytoskeletal components, leading to collective cell movements such as convergent extension; in the absence of Vangl, the cytoskeletal rearrangements that generate epithelial neighbor exchanges are lost, and cells remain in place (Cetera et al., 2018; Goto and Keller, 2002; Kunimoto et al., 2017; Nikolopoulou et al., 2017; Sutherland et al., 2020; Torban et al., 2004). At earlier stages of development, timelapse imaging analysis has revealed the pulmonary mesenchyme to be a highly motile and fluid tissue (Goodwin et al., 2022). Thus, we hypothesized that the mesenchyme might be similarly fluid during sacculation, and that the ability of mesenchymal cells to exchange neighbors might depend on expression of Vangl.

Because of its size, geometry, and need for blood flow and lumenal pressure, timelapse imaging of the late-stage mouse lung remains technically challenging. Therefore, to test the validity of our hypothesis, we turned to mathematical modeling. This approach allowed us to determine the relative roles of mesenchymal fluidity and other known mechanical forces during sacculation. In the model, the epithelial layer consists of chains of cells represented as a curved line, simulating 2D saccules. The epithelial cells experience cell-cell interactions as well as outward movements due to lumenal pressure. The epithelium exhibits a “bending elasticity”, which allows it to change shape in response to forces from the pressure within the lumen (**Fig. 5A**) (Harding, 1997; Kitterman, 1996; Li et al., 2018). We approximated the mesenchyme as a packed field of cells; the ability of the cells within this tissue to exchange neighbors depends on its relative fluidity, which is related to both mesenchymal cell motility (specified by a polarity term) and the forces of cell-cell interactions (specified by a mobility term, which is the inverse of the friction parameter). The model also specifies the (indirect) interactions between the epithelial monolayer and surrounding mesenchymal cells, such that deformation of the epithelium is coupled to rearrangements in the mesenchyme. The initial size and density of the saccules, and the density of the mesenchymal cells, were based on data collected from *E*16.5 wild-type lungs (**Supp. Fig. 4A-F**).

Since previous work has suggested that mechanical forces from fluid pressure are required for sacculation to proceed, we independently varied two parameters in the model: lumenal pressure and mesenchymal fluidity. We found that when lumenal pressure is low, the epithelium fails to expand regardless of the fluidity of the mesenchyme (**Fig. 5Bi-ii, Supp. Fig. 4G-H**). When both lumenal pressure and mesenchymal fluidity are high, the mesenchyme between adjacent epithelial compartments thins and the simulated epithelium expands into saccules (**Fig. 5Biii**). However, when lumenal pressure is high and mesenchymal fluidity is low, the mesenchyme remains thick and expansion of the epithelium is constrained (**Fig. 5Biv**).

**Figure 5.**
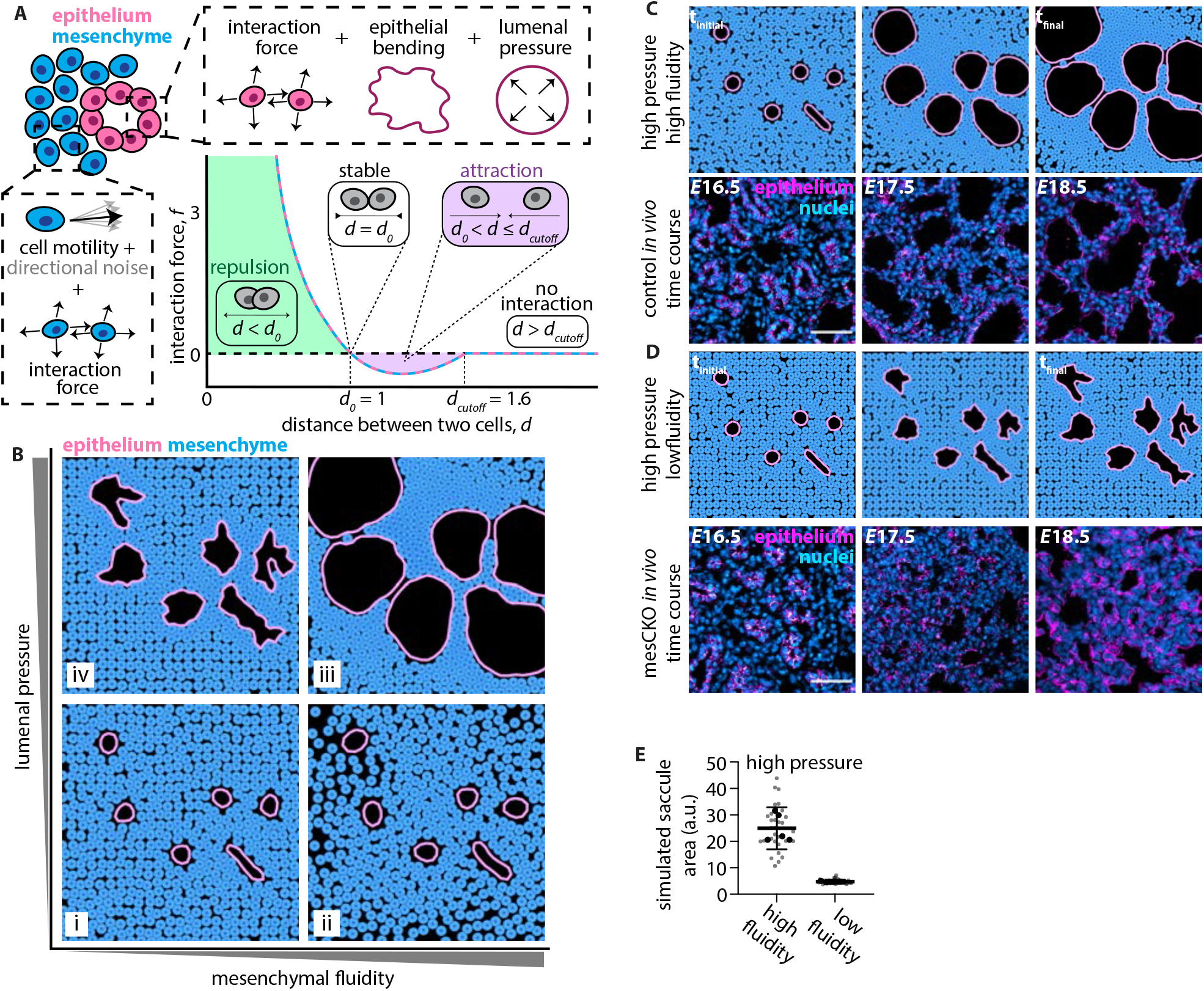
Mathematical modeling predicts that sacculation requires fluidity of the mesenchyme. **A**, Schematic of the parameters used to model the epithelium and mesenchyme. **B**, Phase diagram of final time point of simulations that independently vary lumenal fluid pressure and mesenchymal fluidity. **C**, Snapshots of the timecourse of sacculation in high fluidity, high pressure simulations and in wild-type lungs *in vivo*. Images of *E*16.5-17.5 lung sections immunostained for Ecad (magenta), counterstained with Hoechst (cyan); *E*18.5 lung sections immunostained for RAGE and SPC (both magenta), counterstained with Hoechst (cyan) (scale bars, 50 μm). **D**, Snapshots of the timecourse of sacculation in low fluidity, high pressure simulations and in inducible mesCKO lungs. Images of *E*16.5-17.5 lung sections immunostained for Ecad (magenta), counterstained with Hoechst (cyan); *E*18.5 lung sections immunostained for RAGE and SPC (both magenta), counterstained with Hoechst (cyan) (scale bars, 50 μm). **E**, Quantification of the average saccule area at the final timepoint of high fluidity, high pressure and low fluidity, high pressure simulations.

We found that under high fluidity, high pressure conditions, sacculation *in silico* proceeds similarly to sacculation *in vivo* in wildtype lungs (**Fig. 5C**). The mesenchymal layer between epithelial compartments is thinner at the final timepoint of the simulation than at the initial timepoint, similar to the changes in mesenchymal thickness observed *in vivo* from *E*16.5-18.5. Importantly, simulations in which pressure is high but fluidity is low result in smaller saccules and a thicker mesenchyme, consistent with the morphological defects that we observe when *Vangl1/2* is depleted from the mesenchyme *in vivo* (**Fig. 5D**). Consistently, decreasing the fluidity of the mesenchyme decreases the average saccule area in the simulation, even when the lumenal pressure is high (**Fig. 5E**). Our model therefore predicts that lumenal pressure and a fluid mesenchyme are both essential for sacculation, and that loss of either leads to failure in this morphogenetic process.

### Lineage tracing reveals a fluid mesenchymal cell compartment

Our mathematical model predicts that a fluid mesenchymal compartment is required for epithelial expansion and mesenchymal thinning during sacculation. Our model also predicts that this fluidity results in both active rearrangements of neighboring mesenchymal cells, as well as dispersal of cells during the process of sacculation (**Fig. 6A-C**). To test this prediction experimentally, we used the *Confetti* lineage-tracing system to assess the extent to which mesenchymal cells disperse during sacculation. We generated *Tbx4-rtTA; Tet-O-Cre; Rosa26-Confetti* mice to lineage-label individual mesenchymal cells and follow their progeny through developmental time. In this system, exposure to doxycycline promotes the random expression of one of four fluorophores in a subset of cells within the pulmonary mesenchyme. After injecting a single low dose of doxycycline at *E*14.5, we observed extremely rare groups of GFP^+^ clonal populations at *E*16.5 (**Fig. 6D**). To assess the extent to which mesenchymal cells disperse during sacculation, we conducted an experimental time course from *E*16-18.5 (**Fig. 6D**). We then determined the centroid position of each clonal population and plotted the coordinates of all clonal cells with the centroid as the origin (**Fig. 6E**). While the median number of cells per clone was 4 at both stages, cells in clones at *E*18.5 appeared much further apart than those at *E*16.5, consistent with mesenchymal cell dispersal and a fluid compartment (**Fig. 6E, Supp. Fig. 5**). Measuring the average distance of sister cells from the centroid of each clonal population revealed that cells disperse over developmental time, consistent with the predictions of our mathematical model (**Fig. 6F**). The mesenchyme of the sacculation-stage lung is therefore fluid, and we hypothesize that this fluidity is necessary for epithelial expansion and mesenchymal thinning during sacculation.

**Figure 6.**
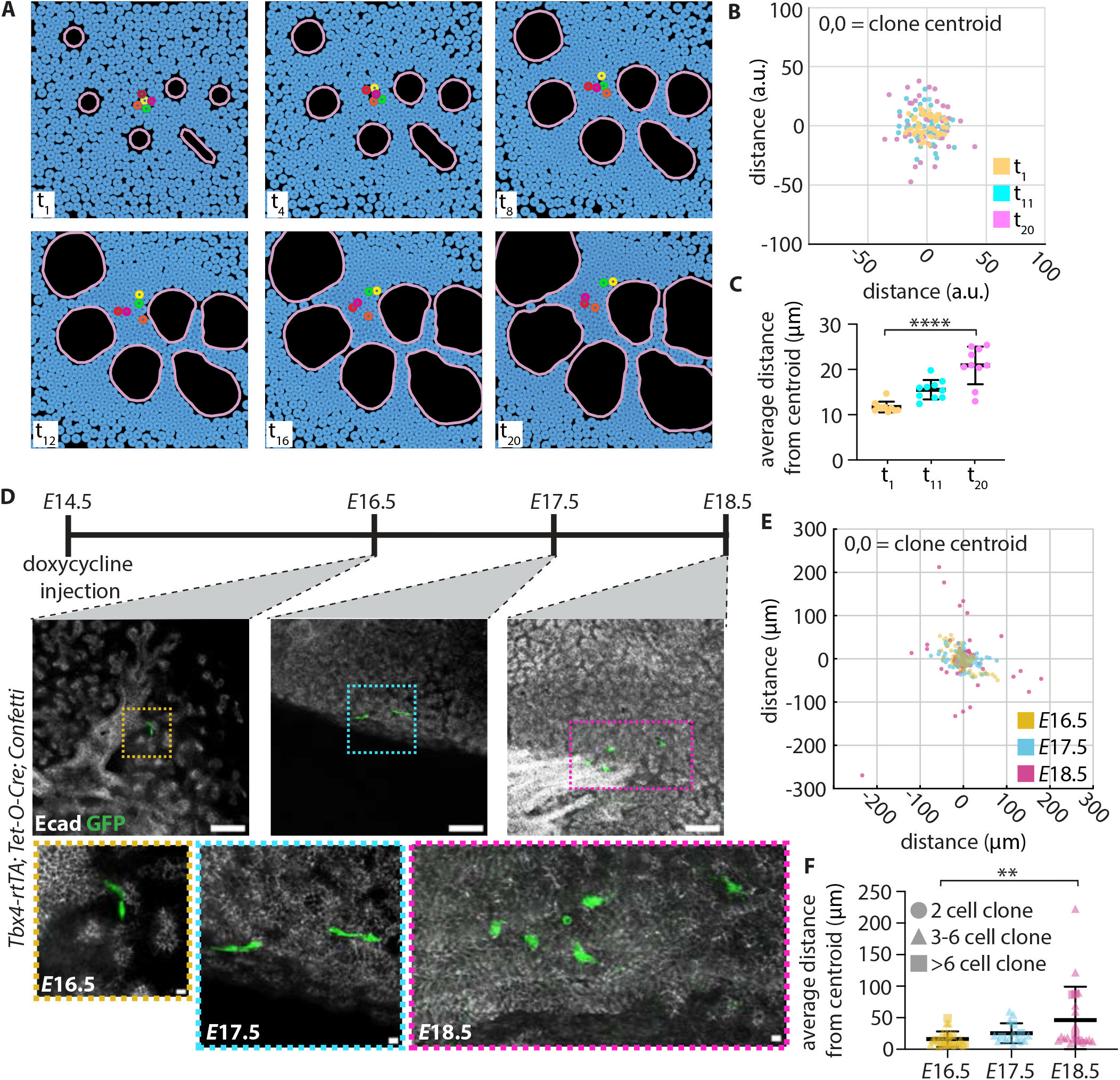
Lineage-tracing reveals a fluid mesenchymal compartment. **A**, Snapshots of timecourse of sacculation in high fluidity, high pressure simulation, with mesenchymal cells color-coded to highlight neighbor exchanges. **B**, Graphical representation of clonal cell positions at t_1_-t_20_ in high fluidity, high pressure simulations; 0,0 is the centroid of each clone. Each dot represents one cell; ten simulations are represented. **C**, Graph indicating the average distance of daughter cells from the centroid of their clonal population at t_1_-t_20_ in high fluidity, high pressure simulations; each dot represents the average distance of all daughter cells from the centroid in one simulation (n=10 simulations, p<0.0001 via unpaired Student’s t-test). **D**, Schematic illustrating experimental protocol for inducing sparse clonal populations, including representative images of clones from 300-μm-thick sections of *Tbx4-rtTA; Tet-O-Cre; Confetti* lungs at *E*16.5-18.5 (scale bars, 100 μm, inset scale bars, 10 μm). **E**, Graphical representation of clonal cell positions at *E*16.5-18.5 in control lungs; 0,0 is the centroid of each clone. Each dot represents one cell. **F**, Graph indicating the average distance of daughter cells from the centroid of their clonal population at *E*16.5-18.5 in control lungs.

### Vangl-mutant mesenchymal cells exhibit an altered cell morphology

Our lineage-tracing data are consistent with our mathematical model, which predicts that the mesenchyme is a fluid compartment during sacculation. We previously found that in early-stage lungs more amenable to timelapse imaging analysis, the fluidity of the mesenchymal compartment correlates with elongated and protrusive cell shapes (Goodwin et al., 2022; Spurlin et al., 2019). We therefore hypothesized that the morphology of mesenchymal cells would be similarly elongated at the sacculation stage in wild-type lungs.

To test this hypothesis, we used a *Tbx4-rtTA; Tet-O-Cre; Confetti* mouse line and examined the shapes of individual mesenchymal cells at *E*17.5. Near the saccular airways, mesenchymal cells exhibited an elongated morphology, often wrapping around the airways (**Fig. 7A**). This elongated morphology is consistent with a more fluid or motile compartment (Goodwin et al., 2022; Spurlin et al., 2019). To determine whether the morphology of mesenchymal cells is altered when *Vangl1/2* are depleted, we introduced the *Confetti* allele into our inducible mesenchymal *Vangl1/2* knockout line (*Tbx4-rtTA; Tet-O-Cre; Vangl1*^*fl/fl*^; *Vangl2*^*fl/fl*^; *Confetti*), where any cell expressing a Confetti fluorophore will also be mutant for *Vangl1/2*. Unlike cells from control embryos, *Vangl1/2*-mutant mesenchymal cells adjacent to distal airways appear less elongated and less protrusive (**Fig. 7B**). To quantitatively describe these changes in cell morphology, we performed automated segmentation of Confetti-labeled mesenchymal cells and analyzed four different shape metrics: aspect ratio, shape factor, circularity, and the ratio of cell area to the area of its convex hull. These cell-shape analyses showed that the *Vangl1/2*-mutant cells significantly differed from control cells in their morphology: mutant cells have an increased circularity, a reduced aspect ratio, a lower shape factor, and a higher ratio of area to convex hull area (**Fig. 7C-F**), morphologies that are consistent with a less fluid or motile state. These data indicate that loss of *Vangl* alters mesenchymal cell morphology, which is consistent with our predictions of a less fluid compartment.

**Figure 7.**
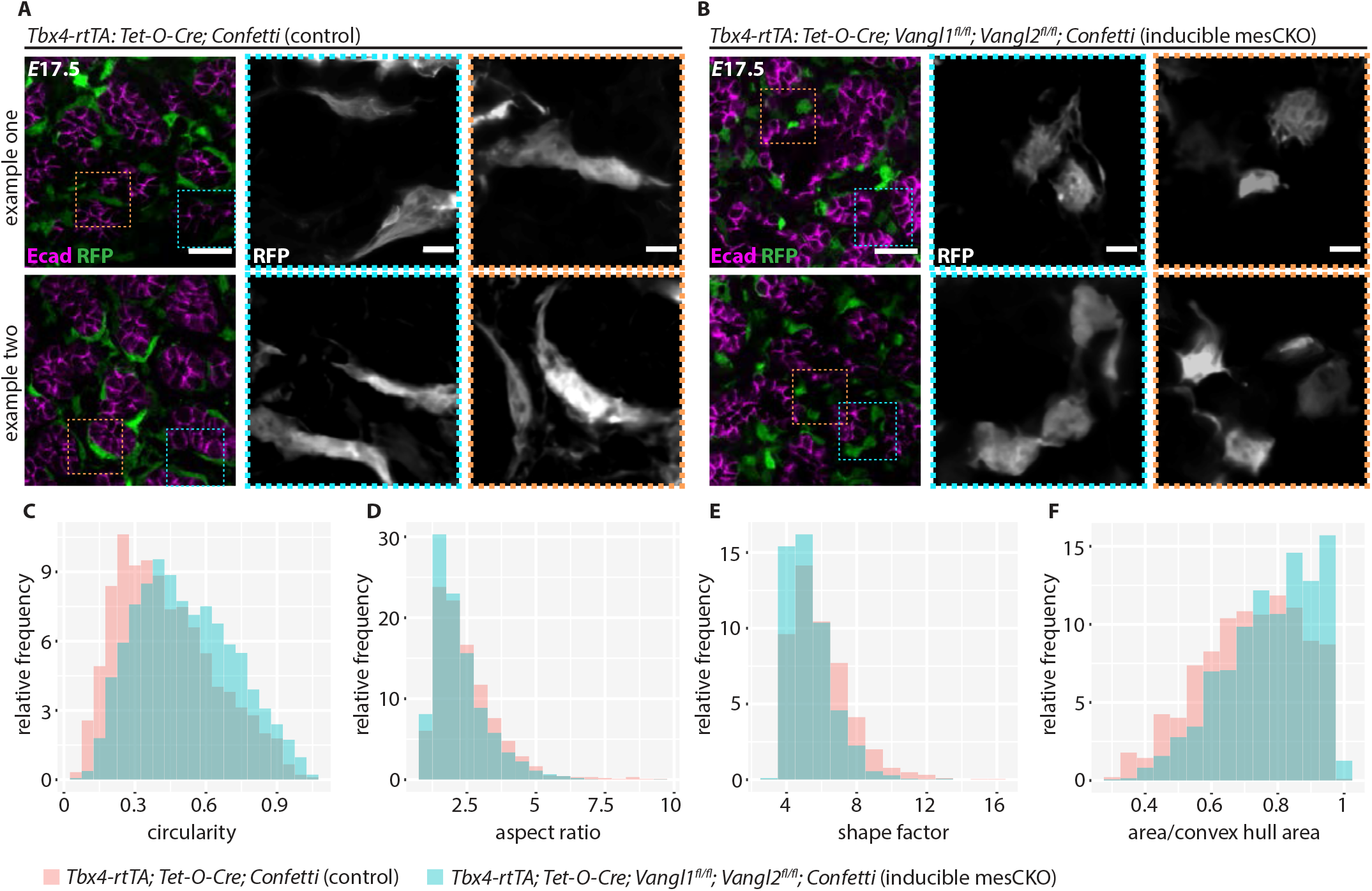
*Vangl2*-mutant mesenchymal cells exhibit altered cell morphologies. **A**, Representative optical sections from two separate *E*17.5 *Tbx4-rtTA; Tet-O-Cre; Confetti* lung lobes. 200-μm-thick sections were immunostained for Ecad (magenta) and imaged for RFP fluorescence (green) (scale bar, 25 μm; inset scale bar, 5 μm). **B**, Representative optical sections from two separate *E*17.5 *Tbx4-rtTA; Tet-O-Cre; Vangl1*^*fl/fl*^; *Vangl2*^*fl/fl*^; *Confetti* lung lobes. 200-μm-thick sections were immunostained for Ecad (magenta) and imaged for RFP fluorescence (green) (scale bar, 25 μm; inset scale bar, 5 μm). **C-F**, Histograms of the area/convex hull area (p < 2.2e^-16^, Wilcoxon rank sum test), circularity (p < 2.2e^-16^, Wilcoxon rank sum test), shape factor (p < 2.2e^-16^, Wilcoxon rank sum test), and aspect ratio (p = 5.953e^-07^, Wilcoxon rank sum test) quantification of mesenchymal cells from *E*17.5 *Tbx4-rtTA; Tet-O-Cre; Confetti* (n = 894 cells from 4 lungs) and *Tbx4-rtTA; Tet-O-Cre; Vangl1*^*fl/fl*^; *Vangl2*^*fl/fl*^; *Confetti* lungs (n=1330 cells from 3 lungs).

## Discussion

Sacculation is an integral part of embryonic development, generating the epithelial surface area required for gas exchange postnatally. However, we still know relatively little about the cellular mechanisms underlying this developmental event. Our data reveal that neither *Vangl1/2* nor *Celsr1*, core members of the PCP complex, are necessary in the pulmonary epithelium for sacculation. Rather, *Vangl1/2* expression in the mesenchyme is essential for mesenchymal thinning and epithelial expansion, key morphogenetic events that drive sacculation. These data are consistent with previous reports of a sacculation defect in the lungs of *Vangl2*^*Lp/Lp*^ mutants (Yates et al., 2010). In contrast to that work, however, we found that most *Celsr1*^*Crsh/Crsh*^ mutant lungs sacculate normally. Because we only observe sacculation defects in lungs of the most phenotypically abnormal *Celsr1*^*Crsh/Crsh*^ embryos, we postulate that the previously reported sacculation defects in *Celsr1*^*Crsh/Crsh*^ embryos are secondary to defects in the vascular system or embryonic demise. These data highlight the need for caution in interpreting genetic manipulations that also affect the gross morphology and viability of the embryo.

Several studies have investigated how alveolar epithelial cell differentiation and the resulting cell-shape changes, particularly the flattening of AEC1s, contribute to sacculation (Frank et al., 2016; Li et al., 2018; Wang et al., 2016). The requirement for both fetal breathing movements (Liggins et al., 1981; Moessinger et al., 1990; Perlman et al., 1976; Tseng et al., 2000; Wigglesworth and Desai, 1982) and the resulting increase in mechanical strain on the epithelium (Huang et al., 2012; Li et al., 2018) has been well established. However, this epithelial-centric focus on sacculation ignores the physical barrier imposed by a thick mesenchymal layer and has left open the question as to whether the mesenchyme might play a more active role in facilitating this stage of morphogenesis. Our study reveals the unexpected finding that *Vangl1/2* expressed specifically in mesenchymal cells plays a role outside of the core PCP complex to drive the formation of saccules. Because Vangl is best known for regulating cytoskeletal organization rather than transcriptional output, we postulate that Vangl1/2 regulates cellular motility to increase mesenchymal fluidity and facilitate active thinning of the mesenchymal cell layer. Our data suggest that in the absence of Wnt5a/Vangl signaling, the forces from lumenal pressure alone are insufficient to thin the mesenchyme. Rather, we predict that Wnt5a/Vangl signaling is upstream of a pathway regulating mesenchymal motility, and that a fluid population of mesenchymal cells is essential for normal sacculation.

Early in vertebrate development, PCP facilitates convergent extension during gastrulation and neural tube closure (Goto and Keller, 2002; Heisenberg et al., 2000; Tada and Smith, 2000; Wallingford et al., 2000). Specifically, PCP pathway proteins are required in mesodermal and neuroectodermal tissues (Butler and Wallingford, 2017; Keller and Sutherland, 2020; Nikolopoulou et al., 2017; Sutherland et al., 2020), which are planar sheets of cells with some mesenchymal properties at these early stages. During organogenesis, the PCP complex has been primarily described in sheets of epithelial cells but is also known to function in planar mesenchymal tissues such as the chondrocytes of the developing limb (Gao et al., 2011). In these planar tissues, PCP protein complexes are asymmetrically aligned along a single axis (**Fig. 1A**).

In contrast, here we show a role for a single PCP component, Vangl1/2, in a complex, nonplanar, 3D mesenchymal tissue that is shaped similarly to a highly perforated block of Swiss cheese. In such a tissue, there is no single plane of cells. In retrospect, it is logical that the entire PCP complex would be dispensable in the pulmonary mesenchyme, as forming a single planar axis of asymmetrically localized protein complexes is physically impossible.

Our study generates exciting new questions that will be investigated in future work. For example, is the fluidity of the mesenchyme a result of directional or random motility°C In our mathematical model, mesenchymal cells move randomly under high pressure, high fluidity conditions, which is sufficient for saccule expansion. Our *in vivo* work suggests that mesenchymal fluidity is regulated by *Vangl* downstream of *Wnt5a*. A recent study investigating the role of *Wnt5a* in tracheal smooth muscle revealed that whereas wildtype tracheal smooth muscle (TSM) cells wrap circumferentially around tracheal epithelium, *Wnt5a*-null TSM cells fail to migrate towards the epithelium despite exhibiting random migration (Kishimoto et al., 2018). That work suggests that *Wnt5a* may be specifically required for directional motion in TSM cells. Perhaps a similar mechanism is at play here, and Vangl regulates not just mesenchymal fluidity but directed motility as well. If so, an important next step will be to understand if Vangl is asymmetrically localized or asymmetrically activated in pulmonary mesenchymal cells.

## Supporting information

High Fluidity High Pressure Simulation

High Fluidity Low Pressure Simulation

Low Fluidity High Pressure Simulation

Low Fluidity Low Pressure Simulation

## Abbreviations

AEC1: alveolar epithelial type I cells
AEC2: alveolar epithelial type II cell;
CKO: conditional knockout
CNT: closed neural tube
ECM: extracellular matrix
Fzd: frizzled
ONT: open neural tube
PCP: planar cell polarity

## Acknowledgements

This work was supported in part by grants from the National Institutes of Health (HL110335, HL118532, HL120142, HL164861, HD099030, HD111539, AR066070, AR068320), the Genetics and Molecular Biology Training Grant of the Molecular Biology Department at Princeton University (T32 GM007388), the Camille & Henry Dreyfus Foundation, and a Faculty Scholars Award from the Howard Hughes Medical Institute. S.V.P was supported in part by a Ruth L. Kirschstein (F31) Fellowship. C.T.Y. was supported in part by a postdoctoral fellowship from the NJCCR. R.S. was supported in part by the NSF Graduate Research Fellowship Program. We thank Dr. Wei Shi (Keck School of Medicine of USC) for generously providing us with the *Tbx4-rtTA; Tet-O-Cre* mice. We would like to thank Dr. Sha Wang and the Microscopy Core Facility at Princeton University, a Nikon Center for Excellence, for their assistance with imaging.

