## Supplemental Figure 1 for "*Vangl* facilitates mesenchymal thinning during lung sacculation independently of *Celsr*"

**Supplemental Figure 1: scRNA-seq of core PCP genes**

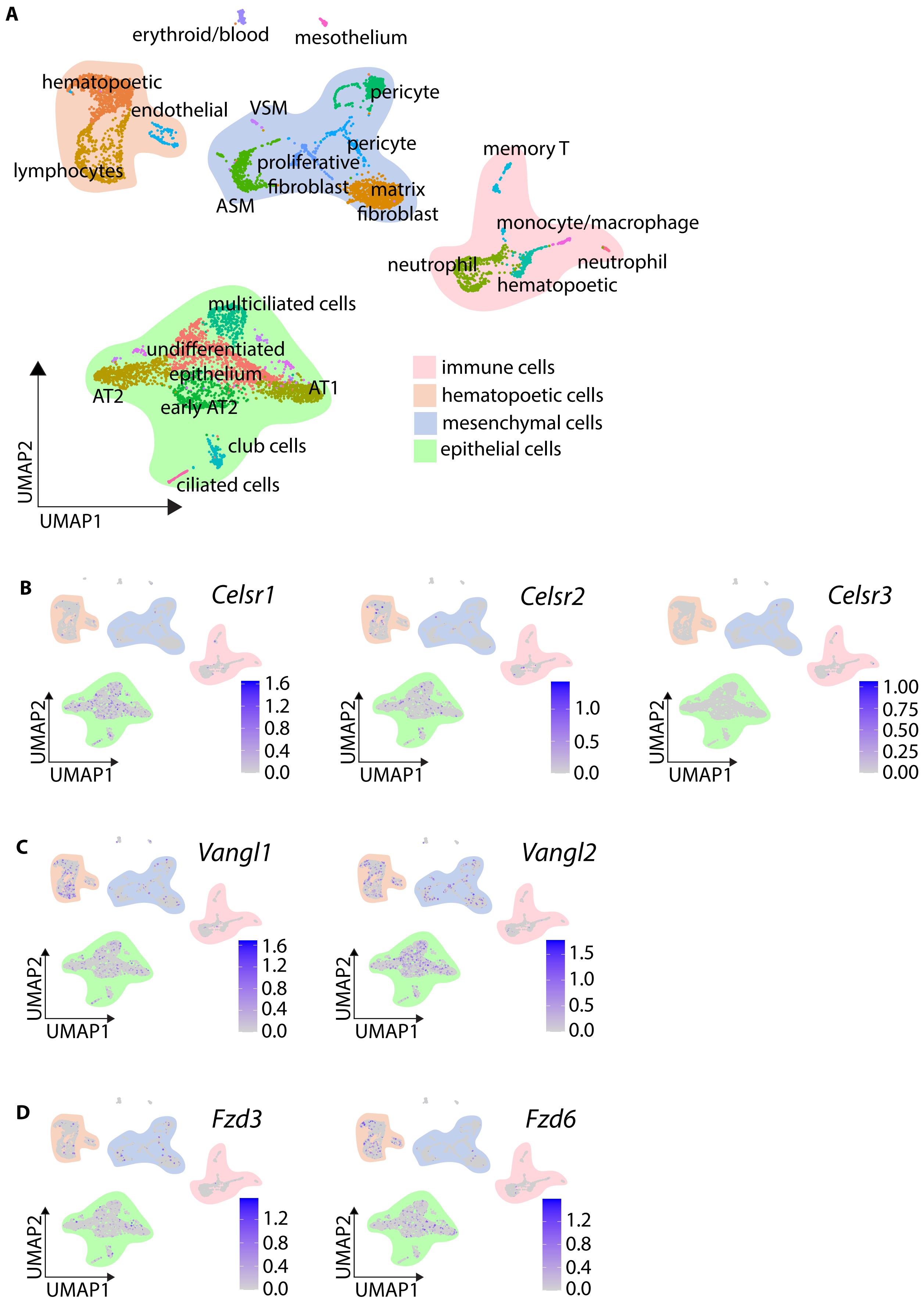

**Supplemental Figure 1. ScRNA-seq of core PCP genes. A,** UMAP projections of *E*17.5 scRNA-seq data; cell-type clusters are annotated based on the five most highly expressed genes within each cluster, and divided into four main categories: epithelial, mesenchymal, immune, or hematopoietic. **B-D,** Expression of each murine core PCP gene is displayed.
