## Supplemental Figure 2 for "*Vangl* facilitates mesenchymal thinning during lung sacculation independently of *Celsr*"

### Supplemental Figure 2: Closed neural tube *Celsr1*$^{Crsh/Crsh}$ embryos exhibit a loss of PCP asymmetry

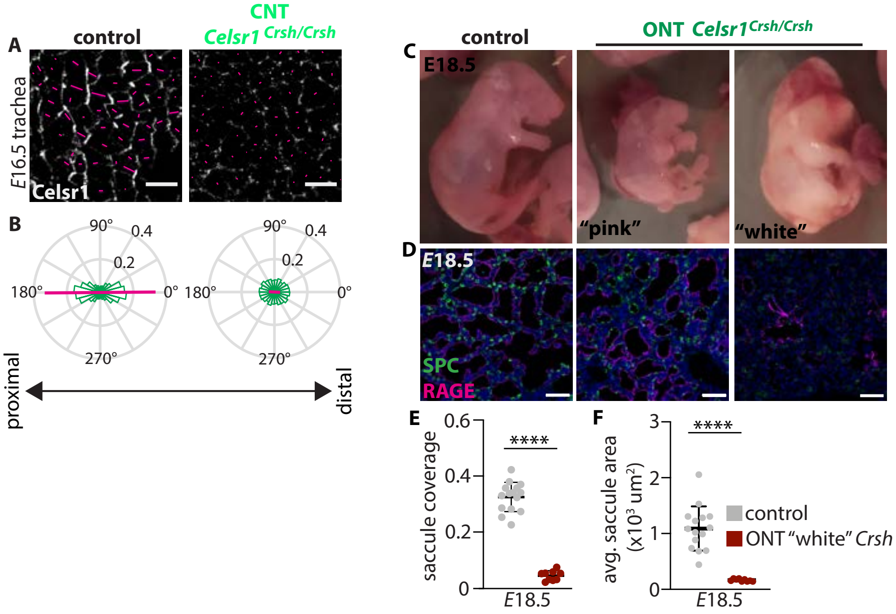

**Supplemental Figure 2. Closed neural tube *Celsr1$^{Crsh/Crsh}$* embryos exhibit a loss of PCP asymmetry. A,** Representative images (scale bars, 10 µm) of immunofluorescence staining for Celsr1 in *E*16.5 tracheas of control and CNT *Celsr1$^{Crsh/Crsh}$* animals quantified in **B**; radial histograms (green) display orientation of Celsr1 asymmetry and magenta line indicates magnitude and direction of average Celsr1; *n*=3 control tracheas, 2970 cells and *n*=3 CNT *Celsr1$^{Crsh/Crsh}$* tracheas, 5121 cells. **C,** Representative images of *E*18.5 control, "pink" ONT, and "white" ONT *Celsr1$^{Crsh/Crsh}$* embryos. **D,** Representative images of sections of distal lung tissue at *E*18.5 in control, "pink" ONT, and "white" ONT *Celsr1$^{Crsh/Crsh}$* embryos; scale bars, 100 µm. **E,** Quantification of the percentage of lung area accounted for by saccules at *E*18.5 in control and "white" ONT *Celsr1$^{Crsh/Crsh}$* lungs (*n*=3 control and *n*=2 mutant lungs, $p$<0.0001 via unpaired Student's t-test); each data point is one lung lobe. **F,** Quantification of average saccular lumenal area at *E*18.5 in control and "white" ONT *Celsr1$^{Crsh/Crsh}$* lungs (*n*=3 control and *n*=2 mutant lungs, $p$<0.0001 via unpaired Student's t-test); each data point is one lung lobe. Shown are mean ± s.d; **** p < 0.0001, as determined by Student's t-test. In all graphs, different shapes represent distinct experimental replicates.
