## Supplemental Figure 3 for "*Vangl* facilitates mesenchymal thinning during lung sacculation independently of *Celsr*"

**Supplemental Figure 3: Compartmental knockouts effectively delete Vangl1/2 in epithelium and mesenchyme.**

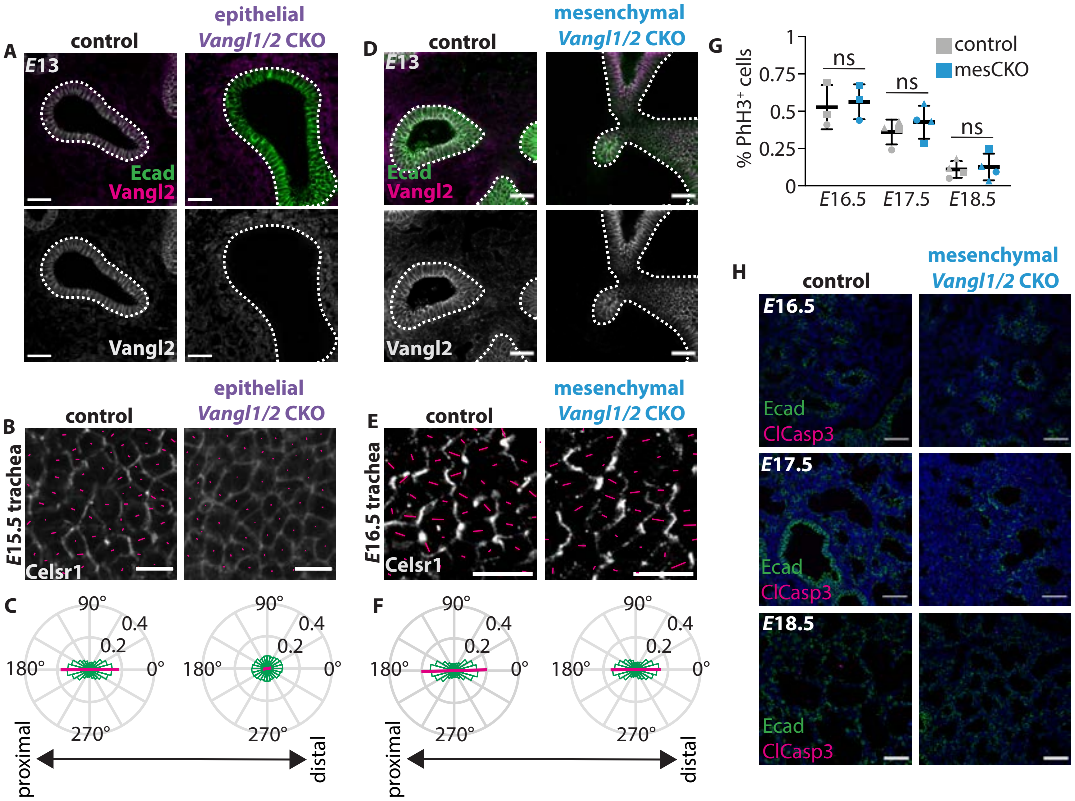

**Supplemental Figure 3. Compartmental knockouts effectively delete *Vangl1/2* in epithelium and mesenchyme. A,** Sections (10-µm-thick) from *E*13 control and epiCKO lungs immunostained for Vangl2 (magenta) and Ecad (green), showing loss of Vangl2 from the epithelial compartment. Dashed lines outline the epithelium; scale bars, 25 µm. **B,** Representative images (scale bars, 10 µm) of immunofluorescence staining for Celsr1 in *E*15.5 tracheas of control and epiCKO animals quantified in **C**; radial histograms (green) display orientation of Celsr1 asymmetry; magenta line indicates magnitude and direction of average Celsr1 polarization; *n*=3 control tracheas, 4288 cells and *n*=3 epiCKO tracheas, 6673 cells. **D,** Sections (10-µm-thick) from *E*13 control and mesCKO lungs immunostained for Vangl2 (magenta) and Ecad (green), showing loss of Vangl2 from the mesenchymal compartment. Dashed lines outline the epithelium; scale bars, 25 µm. **E,** Representative images (scale bars, 10 µm) of immunostaining for Celsr1 in *E*16.5 tracheas of control and mesCKO animals quantified in **F**; radial histograms (green) display orientation of Celsr1 asymmetry; magenta line indicates magnitude and direction of average Celsr1 polarization; *n*=3 control tracheas, 4870 cells and *n*=3 mesCKO tracheas, 5569 cells. **G,** Quantification of cell proliferation in control and mesCKO lungs at *E*16.5 (n=3 control and n=3 mutant, p=0.7536 via unpaired Student's t-test), *E*17.5 (n=4 control and n=4 mutant, p=0.3756 via unpaired Student's t-test), and *E*18.5 (n=4 control and n=4 mutant, p=0.4626 via unpaired Student's t-test). Each data point represents an average of four images from the left lobe of each lung. In all graphs, different shapes represent separate experimental replicates. **H,** Sections (10-µm-thick) from control and mesCKO *E*16.5-18.5 lungs immunostained for Ecad (green) and cleaved caspase-3 (magenta) to label apoptotic cells; scale bars, 25 µm.
