## Supplemental Figure 4 for "*Vangl* facilitates mesenchymal thinning during lung sacculation independently of *Celsr*"

### Supplemental Figure 4: Experimental analysis that defined model parameters

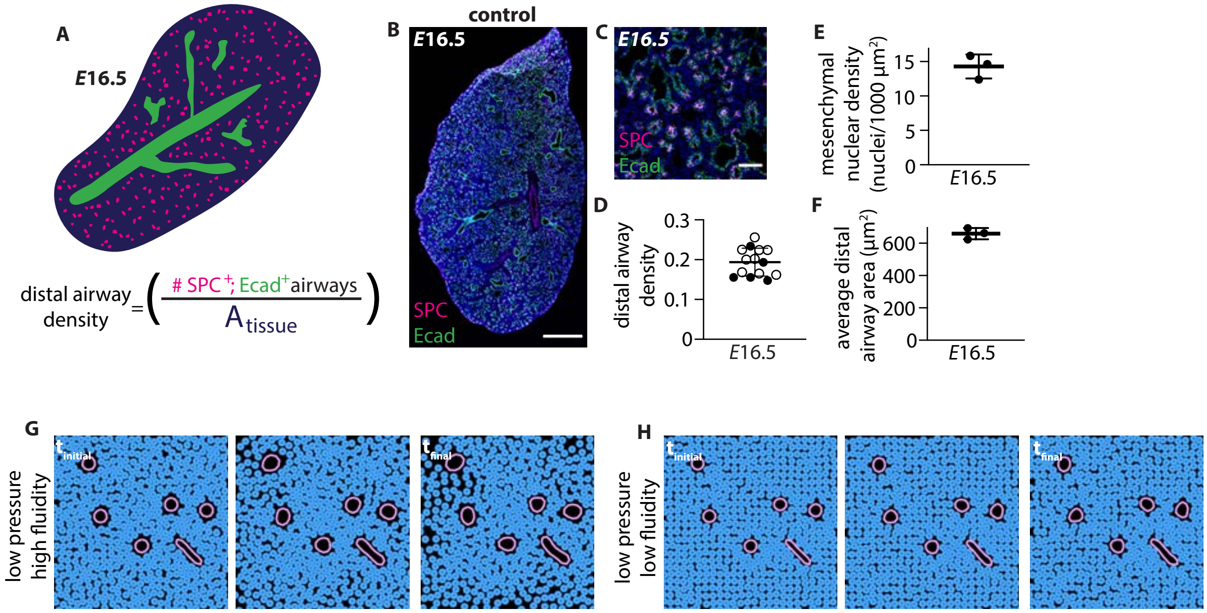

**Supplemental Figure 4. Experimental analysis that defined model parameters. A,** Schematic illustrating metric used to determine density of the distal airways. **B,** Representative tiled images (scale bars, 500 µm) of *E*16.5 lung sections from control lungs immunostained for Ecad (green) and SPC (magenta), counterstained with Hoechst (blue). **C,** Representative images of sections of distal lung tissue at *E*16.5 in control lungs immunostained for Ecad (green) and SPC (magenta), counterstained with Hoechst (blue); scale bars, 50 µm. **D,** Quantification of the density of distal airways per unit area at *E*16.5 in control lungs. **E,** Quantification of mesenchymal cell density in control lungs at *E*16.5. **F,** Average cross-sectional area of distal airways at *E*16.5 in control lungs. **G,** Timecourse of high fluidity, low pressure simulation. **H,** Time course of low fluidity, low pressure simulation.
