## Supplemental Figure 5 for "*Vangl* facilitates mesenchymal thinning during lung sacculation independently of *Celsr*"

**Supplemental Figure 5: Clones from lineage tracing experiments**

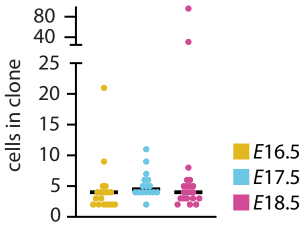

**Supplemental Figure 5. Clones from lineage-tracing experiments.** Quantification of the number of cells per clone in *Tbx4-rtTA; Tet-O-Cre; Confetti* lungs at *E*16.5-18.5; bar on graph represents median. Each dot represents one clonal population.
