## Supplementary figures and images for "*Vangl* facilitates mesenchymal thinning during lung sacculation independently of *Celsr*"

### High Fluidity High Pressure Simulation

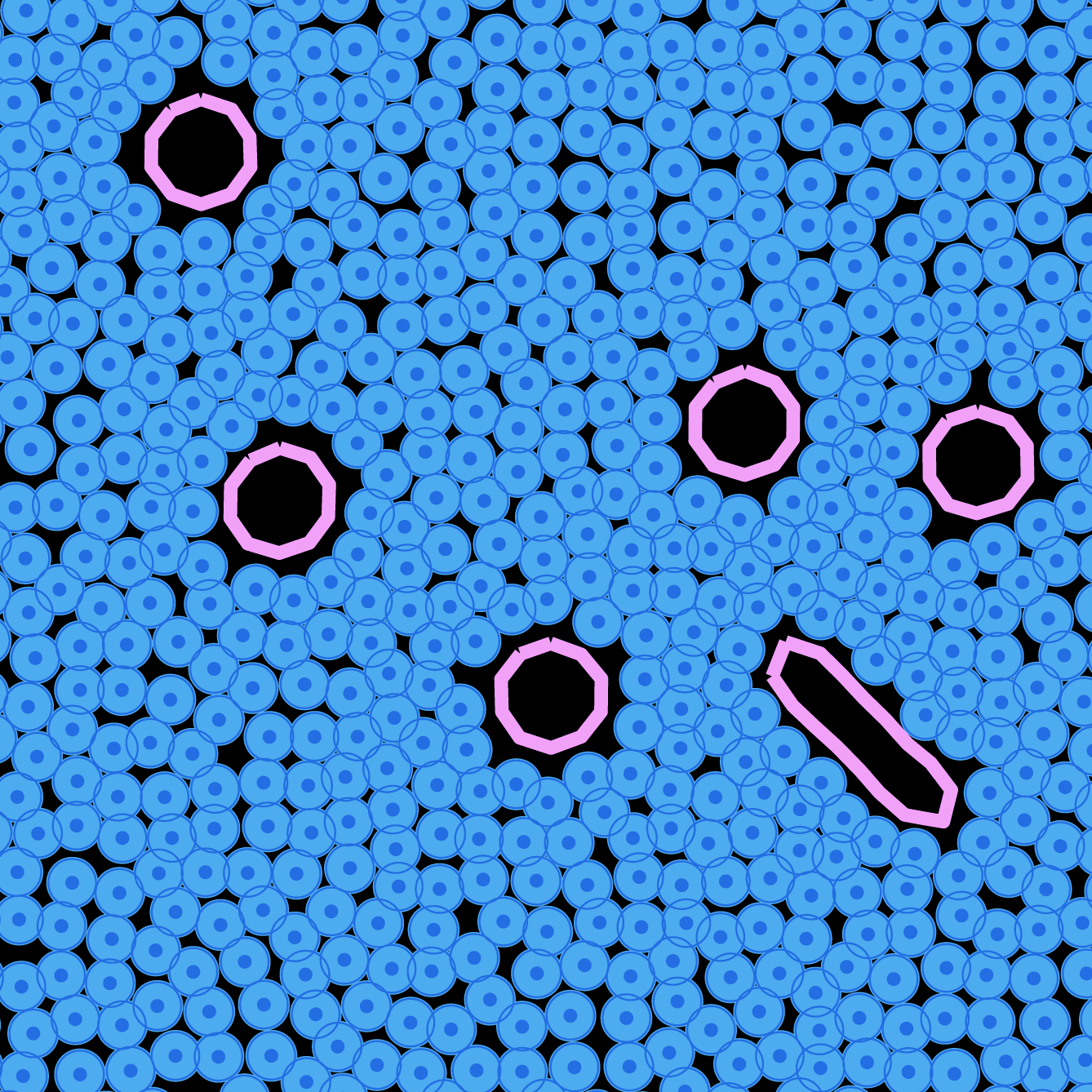

### High Fluidity Low Pressure Simulation

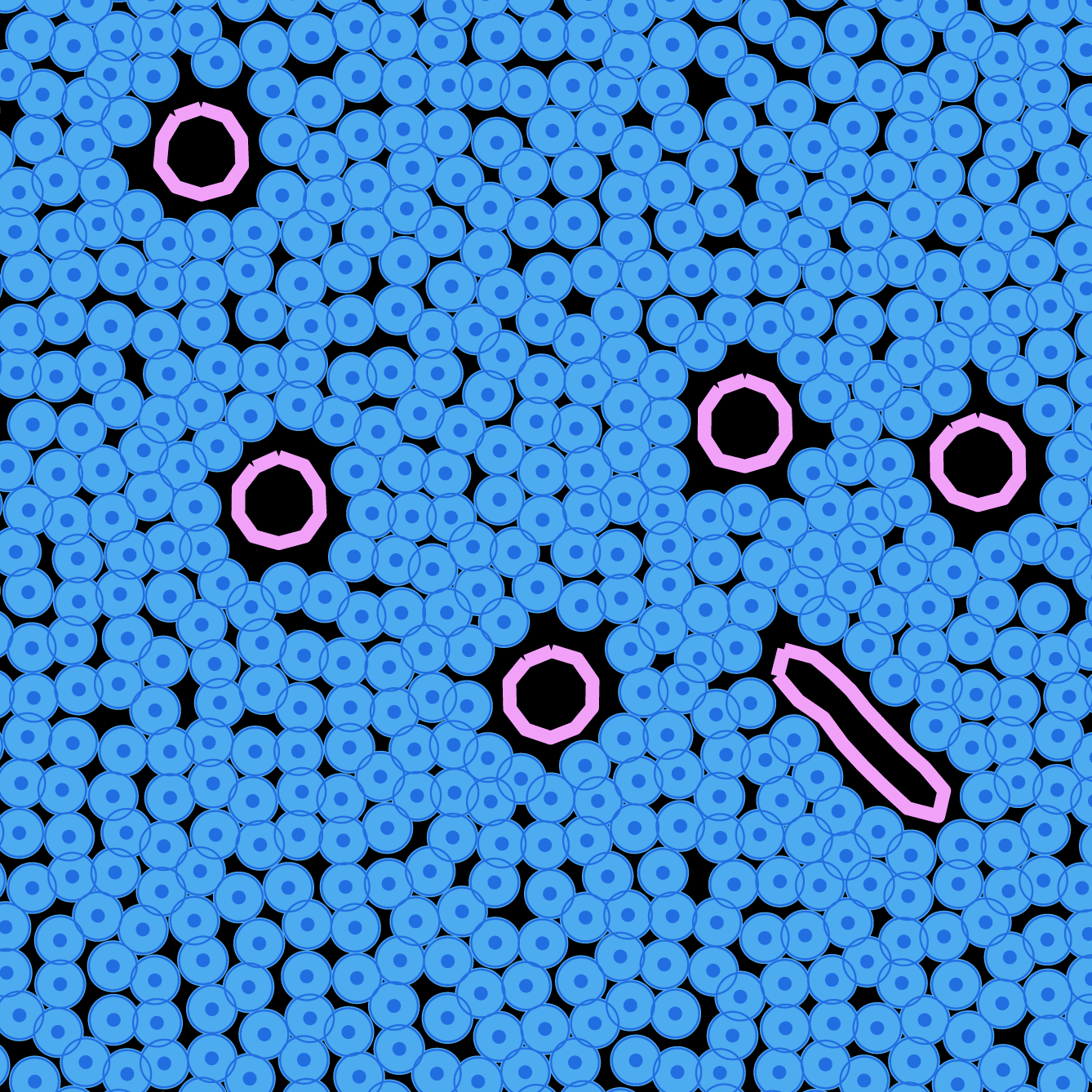

### Low Fluidity High Pressure Simulation

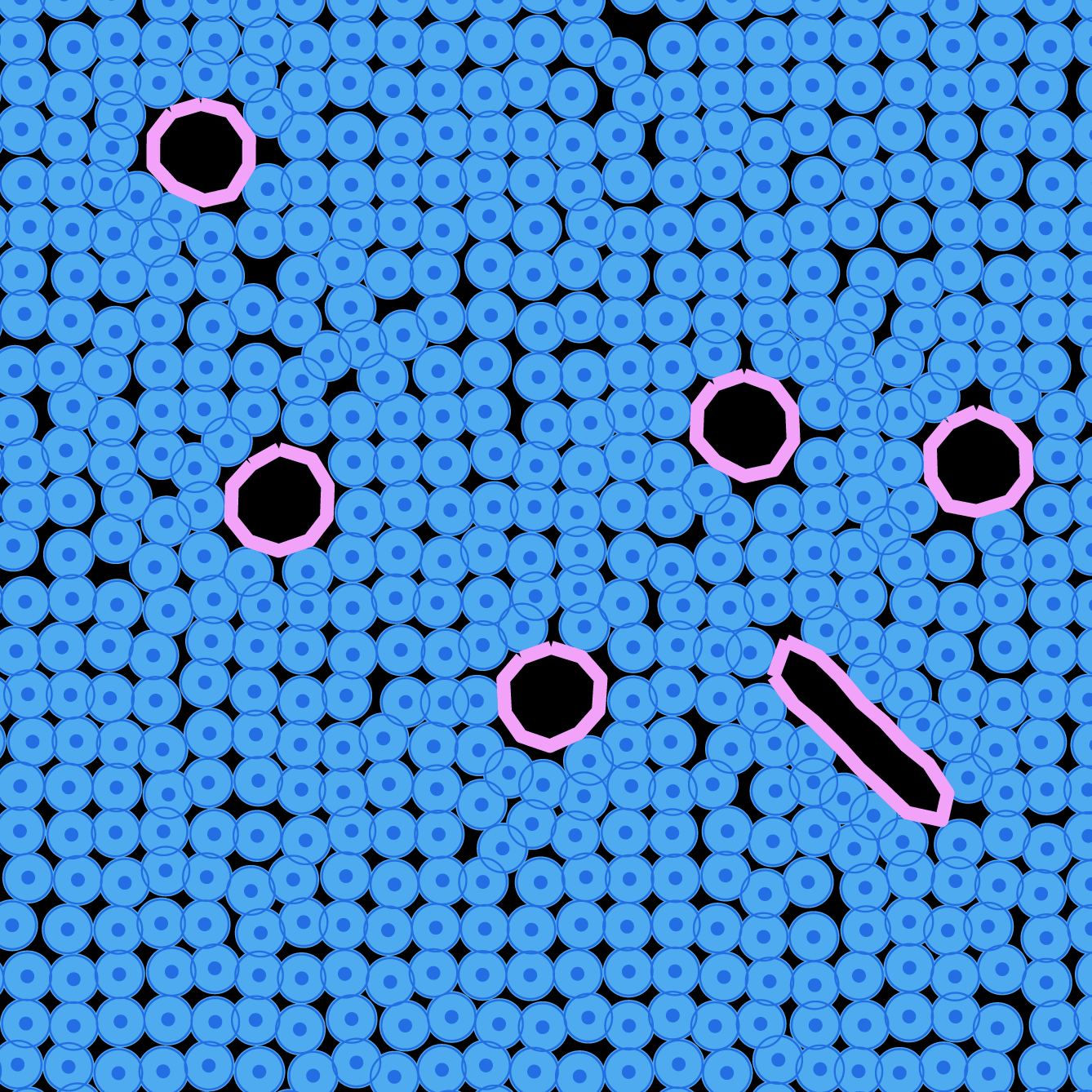

### Low Fluidity Low Pressure Simulation

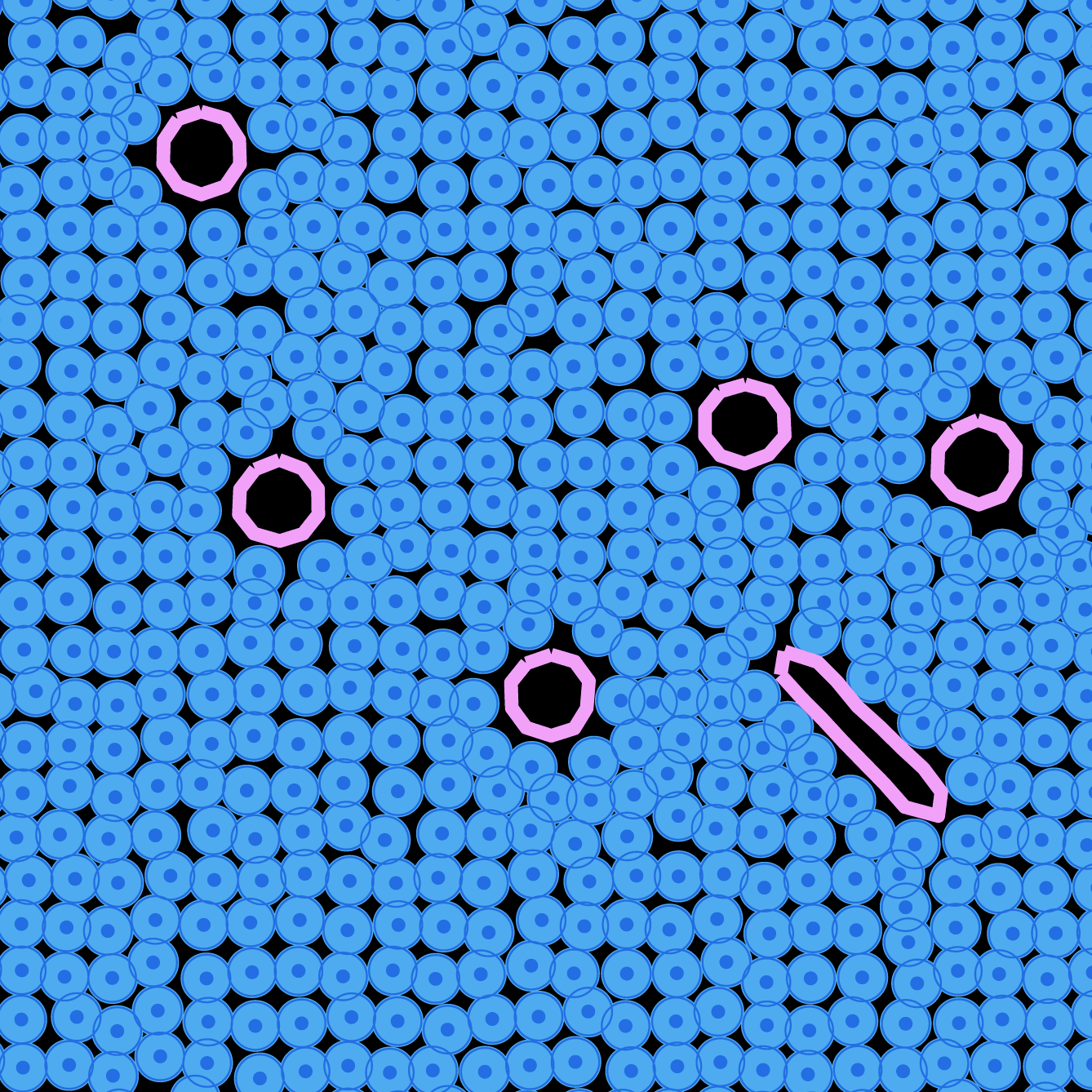
